# A Programmable Toolkit to Dynamically Signal Cells using Peptide Strand Displacement

**DOI:** 10.1101/2021.02.17.431719

**Authors:** Kyle D. Riker, Margaret L. Daly, Micah J. Papanikolas, Tengyue Jian, Stephen J. Klawa, Jacqueline (Yalin) S. Sahin, Dingyuan Liu, Anamika Singh, A. Griffin Miller, Ronit Freeman

## Abstract

The native extracellular matrix communicates and interacts with cells by dynamically displaying signals to control their behavior. Mimicking this dynamic environment *in vitro* is essential in order to unravel how cell-matrix interactions guide cell fate. Here, we present a synthetic platform for the temporal display of cell adhesive signals using coiled-coil peptides. By designing an integrin-engaging coiled-coil pair to have a toehold (unpaired domain), we were able to use a peptide strand displacement reaction to remove the cell cue from the surface. This allowed us to test how the on-demand display of RGDS ligands at variable duration and periodicity of ligand exposure influence cell spreading degree and kinetics. Transient display of α_V_β_3_ selective ligands instructed fibroblast cells to reversibly spread and contract in response to changes in ligand exposure over multiple cycles, exhibiting a universal kinetic response. Also, cells that were triggered to spread and contract repeatedly exhibited greater enrichment of integrins in focal adhesions versus cells cultured on persistent RGDS-displaying surfaces. This dynamic platform will allow us to uncover the molecular code by which cells sense and respond to changes in their environment and will provide insights into ways to program cellular behavior.

## Introduction

The extracellular matrix (ECM) is a dynamic, specialized environment that regulates cell behavior by continually modulating the production, degradation, and remodeling of its components.^1–4^ Engineering of the microenvironment surrounding cells has become a powerful means to study and guide cellular processes such as adhesion, spreading, migration, proliferation or differentiation.^5–7^While great advances have been made in positioning signaling molecules on surfaces to test cellular responses to bound signals and explore the role of their spatial ordering,^8–12^modulating their display in a dynamic, reversible manner has been more challenging to achieve.

Successfully controlling temporal changes in surface-bound cues while maintaining cell viability requires materials that work under physiological pH and temperature, have biocompatible chemistries, induce minimal shear stress, and ideally offer high programmability and orthogonality.^5,13^ This strict set of requirements severely limits the possibilities for synthetically engineered systems. Light responsive,^14,15^ chemically cleavable^16,17^or enzymatically degradable^10–12^ bonds were utilized for epitope display and removal, though most systems thus far offered no or limited reversibility, or operated under potentially harmful conditions to cells. Other efforts engineered electrochemically-responsive surfaces to display or cage signals to promote cell adhesion and differentiation.^18^While this approach enables recycling of the dynamics, it has limited ability for live monitoring of cell behavior. A recent strategy employed to impart dynamic signaling to otherwise inert surfaces involved the use of nucleic acid-peptide hybrids.^19,20^The programmability of DNA allowed the spatiotemporal display of multiple cell instructive signals by harnessing DNA’s ability to undergo strand displacement reactions, in which one strand of DNA is displaced from another by a third strand that binds through unpaired part of the sequence (toehold).21 Inspired by this, we wished to utilize similar dynamic binding of peptides^22^ to regulate cell behavior. Utilizing a pure peptide-based system for both the dynamic switching and cell signaling abilities will not require further chemical modifications and can be readily applicable to soluble or membrane-bound proteins by genetic encoding.^23^

Here we describe the use of strand displacement in coiled-coil dimers to dynamically and reversibly display bioactive signals to cells. By engineering coiled-coil duplexes harboring an adhesion-promoting peptide sequence to be responsive to peptide stimuli, we show that cells can adhere and spread on surfaces reversibly. The use of coiled-coils in cell signalling allows multiple cycles of reversibility with precise temporal control of each cycle. With this capability in hand, one can follow not only cell adhesion, spreading, and subsequent detachment during these dynamic processes, but also dynamic integrin enrichment and reorganization within the cells. This platform presents an attractive model to study the important role of dynamic cell–matrix interactions and provides new avenues for developing rationally designed dynamic cell-instructive materials.

## Results and Discussion

### Synthesis of coiled -coil modified surfaces

In this approach, an input strand (*peptide_I*) (**Figure S1**, for a list of all peptide sequences used, see **Table S1**) consists of a signaling epitope and an α-helical peptide capable of pairing with the α-helical surface strand (*peptide_S*, **Figure S2**) to form a coiled-coil heterodimer as depicted in **Figure 1A**. *Peptide_S* is immobilized onto an alginate-coated glass surface using copper-free click chemistry (**Figures 1B, S3**). As *peptide_I* is longer than *peptide_S* (notation of the heptad repeats of amino acids: H1-H2-H3-H4 compared to H1-H2-H3, respectively), the additional unhybridized peptide segment (H4) serves as a toehold to facilitate interaction and displacement by *peptide_D* (**Figure S4**). To characterize the modified surfaces we performed X-ray photoelectron spectroscopy (XPS) and monitored the elemental composition of the modified surface at every stage of the modification (**Figure S5**). **Figure 1C** shows that the analyzed ratios of C/O and C/N from the XPS measurements were consistent with the predicted monolayer structure for each surface layer.

**Figure 1.**
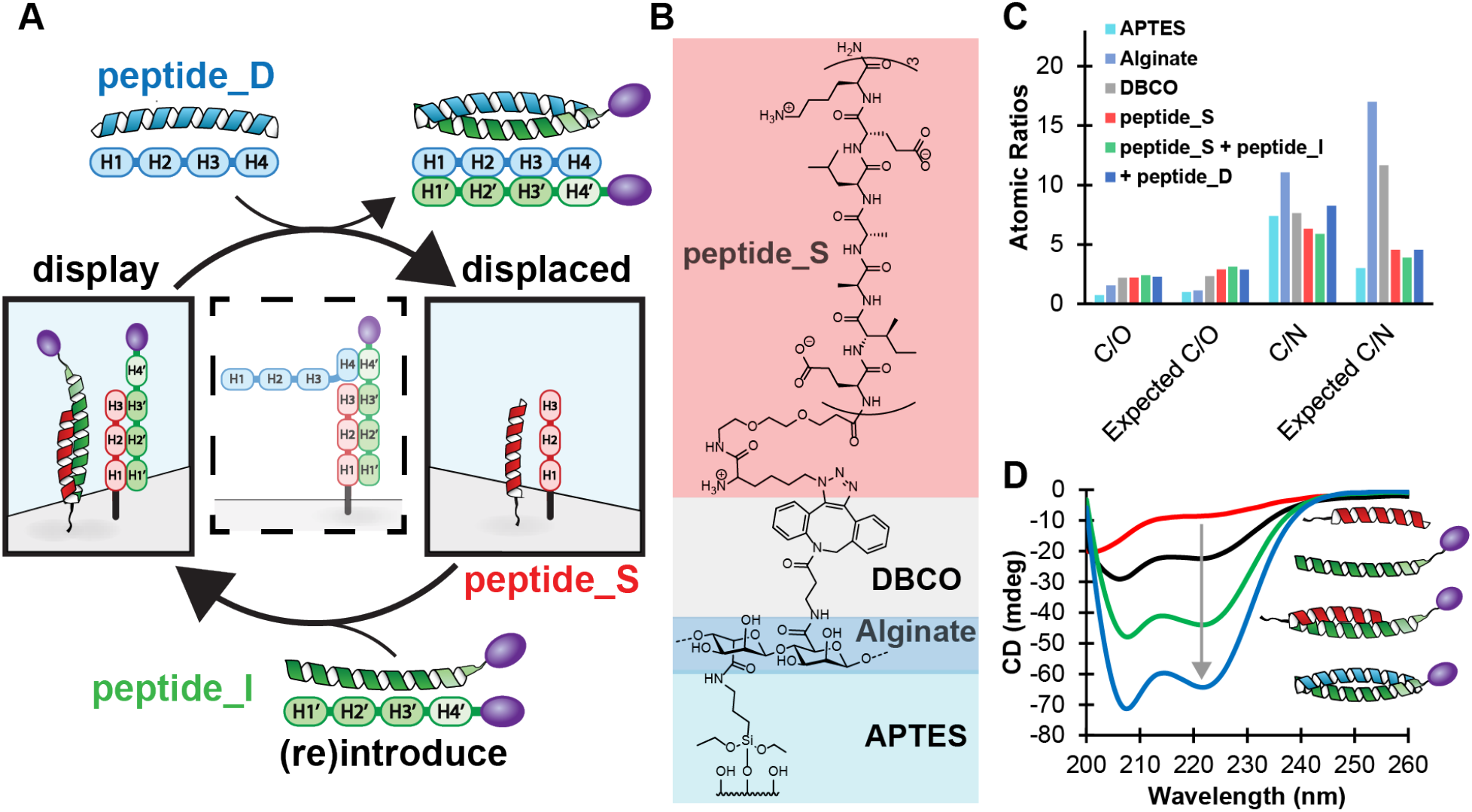
Reversible display of biological signals using coiled-coil modified surfaces. (A) Schematics of coiled-coil strand displacement on surfaces. The middle dashed panel depicts the intermediate state during peptide displacement, where *peptide_D* initially binds *peptide_I* through the toehold. (B) Chemical structure of the surface layers. (C) XPS characterization of atomic composition ratios C/O and C/N at each layer compared to the expected ratios. (D) CD spectra of the various peptides and dimers. Top to bottom: *peptide_S* (red), *peptide_I* (green), *peptide_S* + *peptide_I* (red and green) and *peptide_D* + *peptide_I* (blue and green).

The dynamic reversibility of the coiled-coil duplex formation was assessed using circular dichroism (CD), **Figure 1D**. The individual CD signatures for *peptide_I, peptide_S* and *peptide_D* (**Figure S6**) are typical for α-helices, with a minima at 222 nm and a larger minima at 208 nm. The 1:1 mixture of *peptide_I* and *peptide_S* shows a pronounced minima at 222 nm, indicative of coiled-coil dimer formation. With the addition of *peptide_D*, the 222 nm signal becomes increasingly negative, indicating that the stronger 4-heptad coiled-coil interaction between *peptide_I* and *peptide_D* has formed and displaced *peptide_S*.

### Reversible cell adhesion and morphological analysis

To evaluate the effect of dynamic epitope regulation in coiled-coil modified surfaces on cell behavior, an Arg-Gly-Asp-Ser (RGDS) peptide sequence was incorporated into *peptide_I*. The RGDS epitope was chosen for its ability to engage cell surface integrins (mainly α_5_β_1_and α_V_β_3_) to activate transmembrane signaling pathways that affect cell shape, dynamics, and fate.24,25 We chose to investigate cell adhesion and spreading as a function of display and removal of RGDS using peptide strand displacement, **Figure 2A**. First, to test the accessibility of the epitopes for binding and assess their spatial distribution on the surface, a biotin-containing *peptide_I* (*peptide_I-biotin*, **Figure S7**) was synthesized and incubated with streptavidin-conjugated 20nm gold nanoparticles (AuNPs). Scanning electron microscopy (SEM) images show the AuNPs-labelled peptides were uniformly distributed on the surface, **Figure 2B(i)**, and were successfully removed upon addition of *peptide_D*, **Figure 2B(ii)**. For a quantitative assessment of the peptide loading and average spacing on the surface, we incubated surfaces coated with biotinylated *peptide_I* with horseradish peroxidase (HRP) modified streptavidin, allowing colorimetric quantification of the number of bound *peptide_I*, **Figure 2C(i). Figure 2C(ii)** shows the relationship between the applied concentration of *peptide_I* and its resulting average spacing on the surface (54.8 nm). These results demonstrate that our coiled-coil surfaces can display biological signals with a density similar to natural matrices.^26^

**Figure 2.**
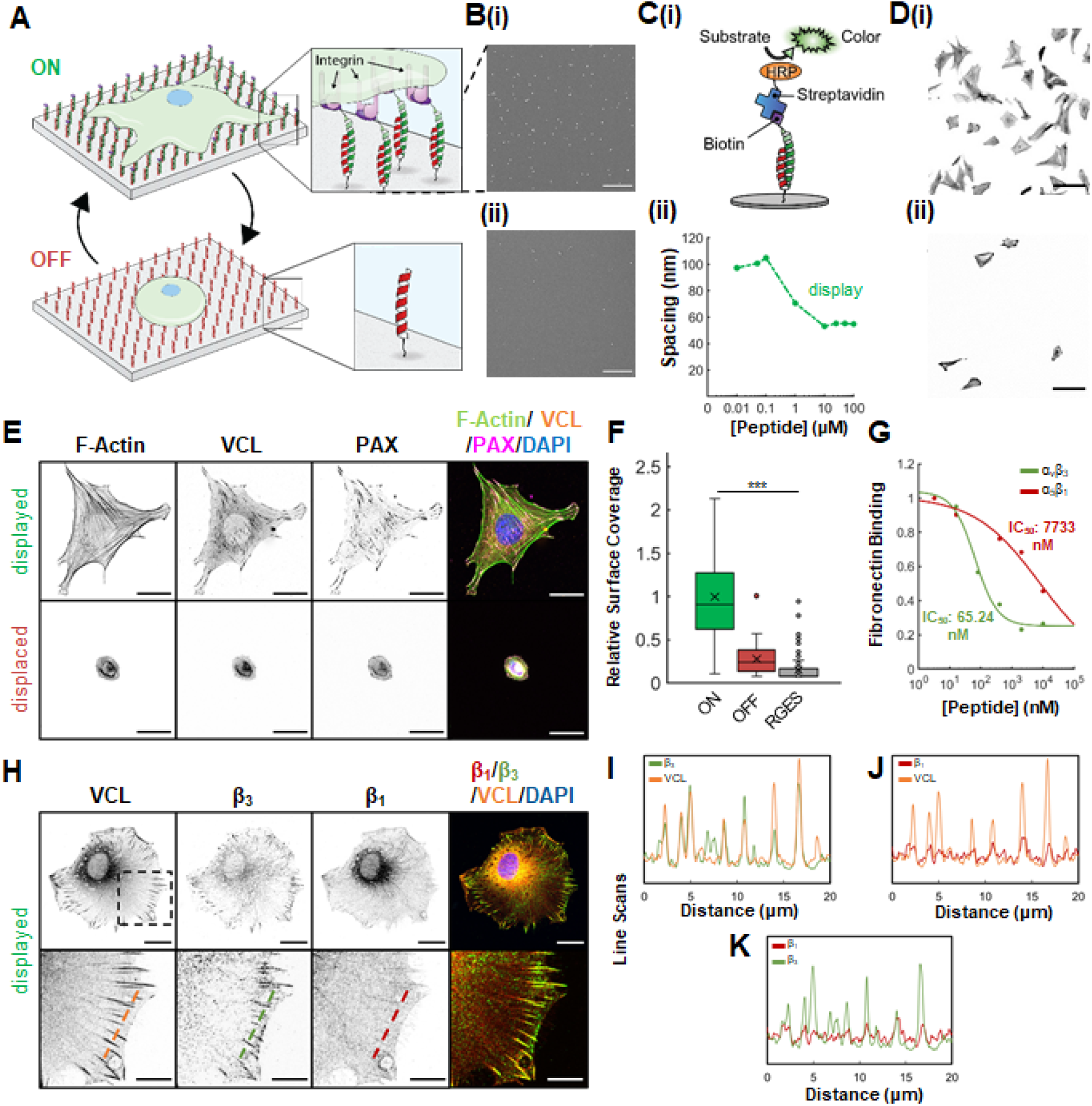
Reversible cell adhesion and spreading on dynamically engineered peptide surfaces. (A) Schematics of reversible display of an integrin-engaging signal to modulate cell adhesion and spreading. (B) SEM images of (i) gold nanoparticles-modified coiled-coil surfaces and the (ii) displaced gold nanoparticles using peptide strand displacement. Scale bars are 200 nm. (C) (i) Schematics and (ii) results of HRP assays to determine the spacing of peptide signals displayed on the surfaces as a function of the concentration of peptide applied. (D) Confocal microscopy images of 3T3 cells plated on a surface before (i) and after (ii) displacement of RGDS from the surface. Scale bars are 100 μm. (E) Confocal microscopy images (split channel and merged) of representative cells on surfaces with displayed and displaced RGDS signal and stained for F-actin (phalloidin: green), vinculin (VCL, orange), paxillin (PAX, magenta) and nuclei (DAPI, blue). Scale bars are 20 μm and insets scale bars are 10 μm. (F) Relative surface coverage of cells on surfaces in the ON and OFF states and control surfaces displaying RGES, quantified from confocal microscopy images. Box and whisker plots show mean (marked by “x”), median (marked by a line) and 25th−75th percentiles. Statistical analysis was performed using an unpaired two-tailed Student’s t-test (***p < 0.001). (G) ELISA results describing the binding of αvβ_3_ and α_5_β_1_ to the coiled-coil signal. Confocal microscopy images (split channel and merge) (H) and line scans (I-K) of cells on surfaces with displayed signal and stained for vinculin (VCL, orange), β_3_ (green), β_1_ (red) and nuclei (DAPI: blue).

Next, we explored if the RGDS-coiled-coil surfaces can induce adhesion, spreading and release of 3T3 fibroblasts cells on demand upon addition of *peptide_D*. **Figure 2D(i)** shows increased fibroblast adhesion and spreading on RGDS-presenting surfaces after 2 hours, while upon addition of the displacer *peptide_D* most cells detached (**Figure 2D(ii)**). SEM images (**Figure S8)** and stained cells for actin, vinculin (VCL) and paxillin (PAX) to visualize the cytoskeleton and focal adhesions (FAs) showed a significant difference in cell size and morphology between the *displayed* (ON) and *displaced* (OFF) states, **Figure 2E**. The RGDS-presenting surfaces had 5x higher levels of surface coverage by adherent cells (**Figure 2F)** compared to surfaces where the signal was displaced. In contrast, plating cells on RGES-displaying (*peptide_I-RGES*) surfaces, **Figure 2F**, or surfaces coated with only *peptide_S* or alginate (**Videos S1-3**) showed poor attachment and no spreading. HEK 293T cells showed limited attachment and no spreading on RGDS surfaces, showcasing that the spreading on RGDS surfaces is mediated by specific integrin activation (**Video S4**).^27,28^

We then explored which integrins the RGDS-coiled-coil surfaces engage with high binding affinity to promote cell adhesion and spreading. As both α_5_β_1_ and α_V_β_3_ integrins bind the RGD amino acid sequence at the FN type III10 repeat,^24,25,29^ we wanted to assess whether our RGDS-appended coiled-coil can selectively engage one integrin over the other. Given the involvement of integrin-mediated adhesion in the regulation of multiple physiological processes^30^ e.g. cell migration, proliferation, survival, and apoptosis) as well as pathological processes (e.g. tumor invasion, metastasis^31^ the development of integrin sub-type selective ligands is highly desirable.

We coated surfaces with either α_5_β_1_ or α_V_β_3_ and incubated them with a mixture of fibronectin (FN) and increasing concentrations of the RGDS-*peptide_I*-*peptide_*S heterodimer (see **Figure S10** for the schematic illustration and details of the assay). FN bound to integrins is detected by a FN-specific primary antibody, while a secondary antibody conjugated with HRP detects bound primary antibody by converting a colorless substrate into a colored product. If the peptide has high affinity for the respective integrin, it will inhibit binding of FN to the surface and no color signal will be detected. By measuring the inhibition of FN binding to immobilized integrins we found that our RGDS-coiled coil is highly α_V_β_3_ selective, with IC_50_ values of 65 nM for α_V_β_3_ and a two orders of magnitude lower affinity (7733 nM) for α_5_β_1_ (**Figure 2G**). This is an interesting result as it shows that compared to a plain RGDS sequence^25^, the high affinity for α_V_β_3_ was not altered by the addition of the coiled-coil (IC50 45 nM vs 65 nM, respectively) yet its selectivity was enhanced by 40x (α_5_β_1_ IC_50_ of 181 nM for RGDS and 7733 nM for *peptide_I*). To visualize integrin engagement, we immunostained fibroblasts cells that were cultured on *peptide_I*-displayed surfaces for 2 hours, **Figure 2H**. Peripheral FAs were observed with highly clustered β_3_ and low β_1_, confirming the selectivity of RGDS-*peptide_I* to α_V_β_3_, **Figure 2I-K**. Focal adhesions were not observed in cells on control surfaces without RGDS-*peptide_I* (**Figure S11**).

### Cell response to temporal variations in RGDS display

As our peptide-based platform allows us to change ligand presentation at will, we wanted to study how cells respond to changes in the microenvironment. To gain insight regarding the possible role of temporal variations in RGDS display on integrin engagement and focal adhesion formation, we designed a set of experiments in which fibroblasts were cultivated on RGDS-displaying surfaces for selected time intervals, after which RGDS was displaced by addition of *peptide_D* (**Figure 3A**). We chose time frames for initial stages of cell spreading (0.5, 1, 2, and 3 hr post-seeding) when contributions from cell-secreted matrix and substrate remodeling are minimal. As shown in **Figures 3B-E and S12**, cells quickly reacted to longer durations of ligand display by increasing area and enhancing number and aspect ratio of their focal adhesions. Interestingly, changes in ligand display via peptide strand displacement promoted a similar decrease in cell area and a reduced number of focal adhesions, regardless of initial adhesion duration (30 min - 2 hr).

**Figure 3.**
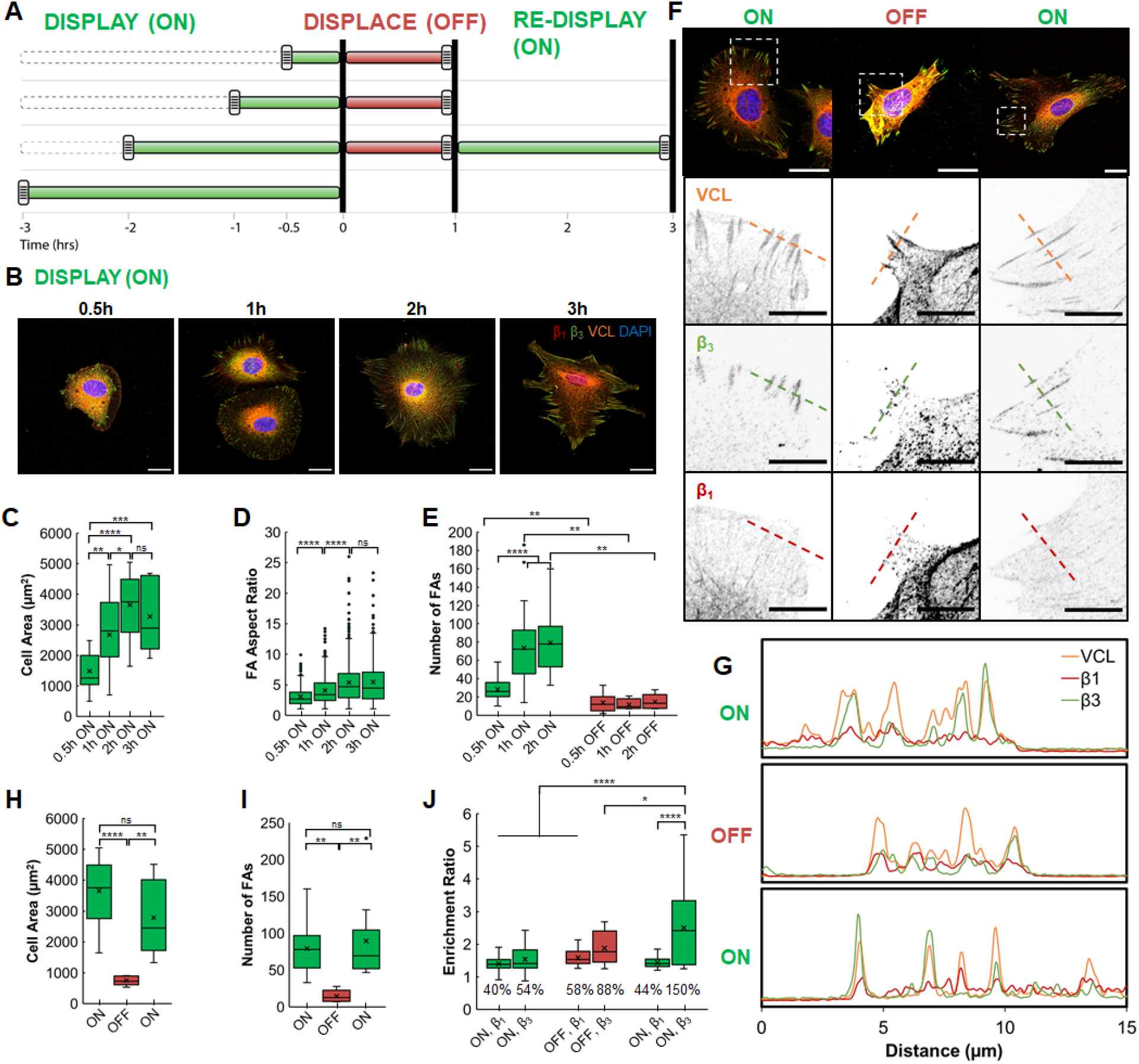
Cell response to temporal variations in ligand display. (A) Scheme representing different temporal patterns of ligand exposure. (B) Confocal microscopy images of cells cultured on RGDS surfaces for various durations. Scale bars are 20 μm. (C) Cell area at different ON time points. (D) Aspect ratio of focal adhesions at various O time points. (E) Number of FAs found per cell. (F) Confocal microscopy images of cells across an ON-OFF-ON cycle. Scale bars on insets are 10 μm (G) Line scans over insets show degree of co-localization of integrins in FAs. Cell area (H) number of FAs (I), and integrin enrichment (J) before, during, and after the displacement step. Cells were stained for β_1_ (red), β_3_ (green), Vinculin (orange), and DAPI (blue). For all panels, addition of *peptide_I*: ON, green; addition of *peptide_D*: OFF, red. All box and whisker plots show mean (marked by “x”), median (marked by a line) and 25th−75th percentiles. Statistical analysis was performed using unpaired two-tailed Student’s t tests in panels (C,E-J). Statistical analysis was done using a Kruskal-Wallis test in panel D. Statistical significance denoted: n.s. p>0.05, * p < 0.05, **p < 0.01, ***p < 0.001, ****p < 0.0001.

Next, we wanted to explore how cells that were previously subjected to display and displace steps respond to re-display of the RGDS ligand. **Figure 3F** shows immunostained cells with visualized focal adhesions and integrins at the various cycle steps of ligand display and removal (ON-OFF-ON). FAs of cells in both display and re-display stages were abundant in α_V_β_3_ integrins, while co-clustering of α_V_β_3_and α_5_β_1_ was not observed (**Figure 3G**). Cells that were re-subjected to RGDS spread back to areas comparable to their initial spreading period (**Figure 3H**), with similar number of focal adhesions (**Figure 3I**). Surprisingly, ligand re-display prompted a highly enhanced enrichment of β_3_ integrins in the focal adhesions, in comparison to the first display step (150% versus 54%), as quantified by the ratio of antibody staining intensity inside adhesion clusters to the intensity in a surrounding region of interest (for details on enrichment ratio calculation see **Figure S13**).^29^ By contrast, the enrichment ratio for α_5_β_1_ integrins was significantly lower than that for α_V_β_3_ integrins, and was not sensitive to the changes in RGDS display, in line with an α_V_β3 -selective surface. (**Figure 3J**). This result highlights the important role of temporal control of integrin engagement in recognizing, perceiving and responding to temporal regulation of integrin-engaging ligands, which in turn enables cells to sense and adapt to their microenvironment. Unraveling how cells respond to these dynamic changes will further our understanding of processes involving extensive remodeling such as embryogenesis, angiogenesis, and cancer progression._32_

To gain further insight into the effect of temporal presentation of cell-adhesive signals, we cultured fibroblasts on *peptide_S* modified surfaces while alternating the adhesive RGDS signal between displayed and displaced states over multiple cycles (**Figure 4**). Live cell imaging showed that cells are able to reversibly spread and retract in response to temporal variations in RGDS display (**Figure 4A and Video S5**). Remarkably, deriving the kinetic curve for each individual cell unraveled universal kinetics of spreading and contracting across the entire cell population (**Figures 4B, Videos S6, S7**). Moreover, repetitive display steps exhibited similar spreading rates (∼43 μm^2^/min) and faster than on aptes or glass surfaces (**Figure S14**). In addition, cell contracting rates in response to a change in ligand display were much faster than the spreading rates (∼95 μm^2^/min) (**Figure 4C**). Interestingly, the cell contraction time frame was similar to the kinetics of peptide strand displacement (**Figure S15**) highlighting the dynamic ‘sense and response’ of integrins to matrix changes in ligand display. To monitor the assembly and disassembly of FAs in response to a cycled display of RGDS, we fixed the cells at various time points and stained them to visualize the cytoskeleton and focal adhesions. **Figure 4D** shows cells with assembled and disassembled FAs in response to the display and displacement of the RGDS signal, respectively. Quantification of the number of FAs validates the microscopy observations showing an enhanced number of FAs on RGDS surfaces and a significantly reduced number of FAs when RGDS is displaced (**Figure 4E**). Moreover, the linear relationship between the number of FAs per cell and cell area (**Figure 4F**) resulted in significant differences between the ‘on’ (display) and ‘off’ (displace) states. It should be noted that cells attached, detached and reattached reversibly, and remained viable throughout all display and displace cycles (**Figures S16**,**S17)**. Taken together, these results demonstrate how temporal display of a surface-bound cell adhesive signal elicits control of cell spreading, contraction, detachment and reattachment, and how fine modulation over multiple cycles exerts control of focal adhesion dynamics and integrin engagement. This opens up the fascinating possibility that, similarly to how soluble biochemical signals elicit different cellular responses according to their dose and their temporal presentation,^4^ there might be an optimal temporal presentation of bound signals to elicit distinct cell responses.

**Figure 4.**
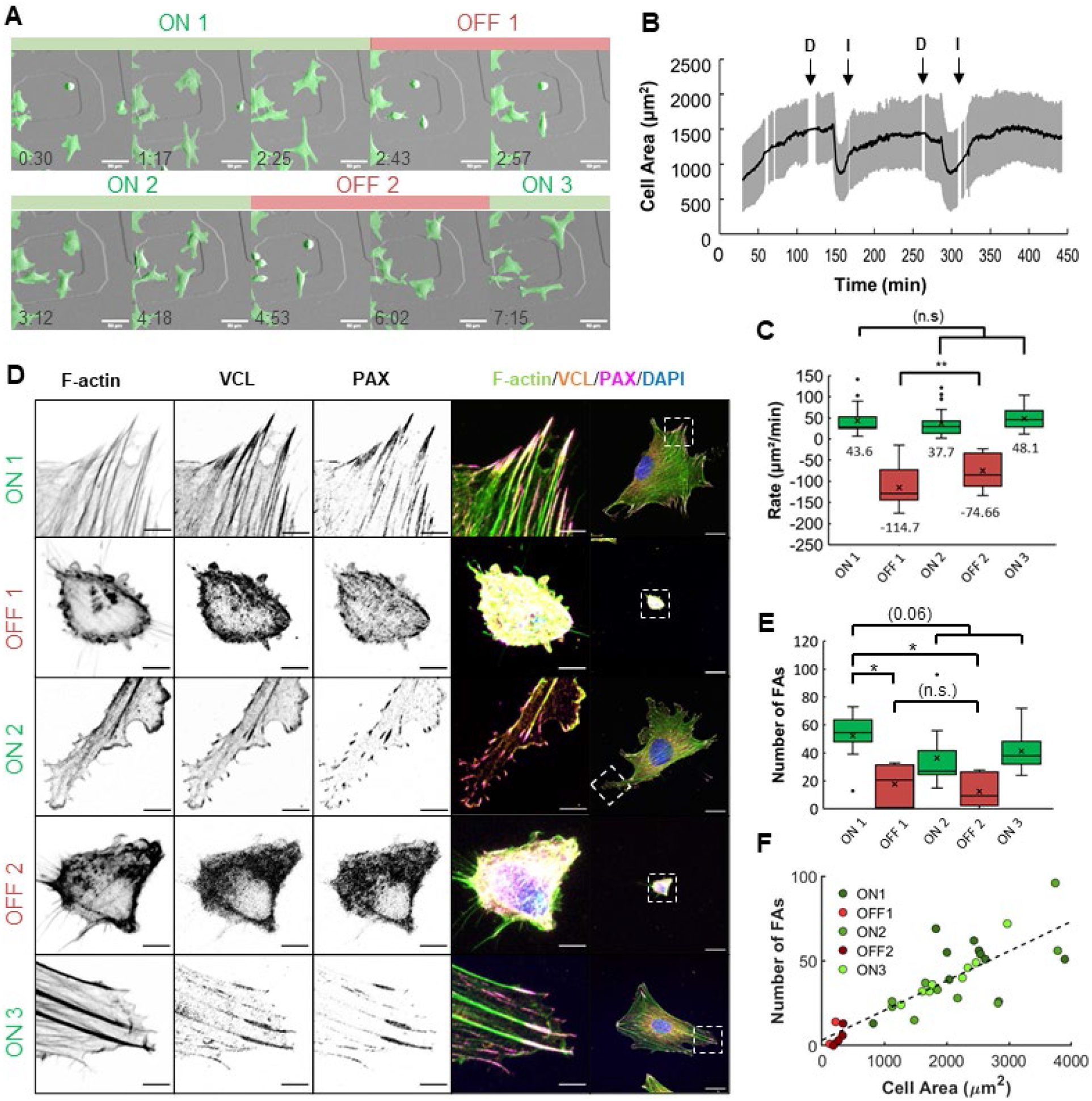
Cell response to alternating display of RGDS over multiple cycles. (A) DIC snapshots of videos tracking the cell spreading and contracting behavior. The green mask outlines the cells in the frame, enabling live single-cell tracking. (B) Average (black line) and standard deviation (gray area) of cell area measured over time from the DIC video (n = 26). (C) Initial rates of spreading (green) and contracting (red) (n = 24). (D) Confocal microscopy images (split channel and merged) of cells stained for F-actin (phalloidin: green), vinculin (VCL, orange), paxillin (PAX, magenta) and nuclei (DAPI, blue). Scale bars are 20 μm, and inset scale bars are 5 μm. (E) Number of focal adhesions per cell. (F) Number of focal adhesions as a function of cell area (correlation coefficient = 0.68). For all panels, addition of *peptide_I*: ON, green; addition of *peptide_D*: OFF, red. All box and whisker plots show mean (marked by “x”), median (marked by a line) and 25th−75th percentiles. Statistical analysis was performed using unpaired two-tailed Student’s t tests (n.s. p>0.05, * p < 0.05, **p < 0.01, ***p < 0.001, ****p < 0.0001).

## Conclusions

We demonstrate a novel method for dynamic substrate engineering using strand displacement with coiled-coil peptides. This dynamic platform enabled us to learn how cells respond to temporal variations of adhesive surface-bound signals. Specifically, we were able to instruct fibroblasts to adhere and spread on surfaces reversibly through integrin-mediated processes over multiple cycles, and demonstrate a temporal pattern of ligand display that led to a uniform and reproducible response by the cells. Importantly, the incorporation of bioactive coiled-coil pairs to generate matrices with dynamic control over bioactivity and cell behavior is not limited to surfaces or to the RGDS sequence, and can be extended to hydrogels or other scaffolds. Moreover, the ability to design multiple peptide handles with variable toeholds (unpaired domains) will enable us to control multiple cellular cues in an orthogonal manner to elucidate their roles and interrelationships in regulating cell decision-making. Finally, the use of peptide triggers has exciting promise for extending our approaches to the *in vivo* environment. Dynamic tuning of cues *in vivo* will have implications for examining a broad array of diseases. We believe this type of dynamic temporal control of cell-material interactions will advance mechanistic understanding of matrix cell biology as well as provide prototypes for tissue engineering and strategies for therapeutics.

## Materials and Methods

### Materials

All purchased chemicals were used without further purification. Rink amide MBHA resin and Fmoc-Lys(azide)-OH were purchased from Chem-Impex. Fmoc-(PEG)2-OH was purchased from PurePEG. Alginate, Fmoc-protected amino acids, HBTU, PyBOP, Oxyma pure, DIC, TFA, TIPS, EDC and NHS were purchased from Sigma-Aldrich. Fmoc-Lys(Alloc)-OH was purchased from P3 BioSystems. Dibenzocyclooctyne-amine (DBCO-amine) was purchased from Sigma-Aldrich, then dissolved in dry DMSO to a final concentration of 100 mM and stored at -20 °C.

## Methods

### Peptide synthesis and purification

All coiled-coil peptides were synthesized using automated standard fluoren-9-ylmethoxycarbonyl (Fmoc) solid-phase peptide synthesis method (Liberty Blue, CEM) on rink amide MBHA resin (100-200 mesh, 0.77 mmol/g). Peptides were cleaved from the resin using a solution of 95% trifluoroacetic acid (TFA), 2.5% triisopropylsilane (TIPS) and 2.5% dH_2_0. The acid was evaporated and the crude peptide was purified using reverse-phase HPLC (Shimadzu UFLC, Ultra C18 5 uM, 100 x 10 mm column) with a gradient of 0.1% TFA in water (Solvent A) and acetonitrile (Solvent B) over 50 minutes. Purified peptides were lyophilized and stored at -20° C. Purity was confirmed by electrospray ionization mass spectrometry and HPLC.

*Peptide_I-biotin* was synthesized using automated standard fluoren-9-ylmethoxycarbonyl (Fmoc) solid-phase peptide synthesis method. The alloc group was deprotected with the addition of 0.1 equiv. of tetrakis(triphenylphosphine)palladium(0) and 20 equiv. of phenylsilane in 6 ml DCM. After 12 hours under room temperature, the resin was washed and added with biotin (2 equiv), DIPEA, HBTU and PyBOP. Solution was stirred for another 12 hours, and the resin was washed and purified to get the final product.

### Preparation and characterization of peptide functionalized surfaces

Round coverslips (12-mm diameter, Carolina Biological Supplies) were marked on one side using a laser cutter in the BeAM makerspace facility of University of North Carolina at Chapel Hill to keep track of the side to be modified. Coverslips were cleaned using a 2% micro-90 solution in water for 30 minutes at 60 °C. After rinsing 6 times with water and 2 times with ethanol, the coverslips were dried and plasma etched for 5 minutes.

The etched coverslips were coated with (3-Aminopropyl)triethoxysilane (APTES, Sigma) by vapor phase deposition. Briefly, 50 µL of 2% APTES in ethanol was added to a small aluminum foil “boat” placed inside a petri dish. Coverslips were placed in the petri dish, and the lid was replaced with a weight on top to prevent APTES vapor from escaping. Coverslips were modified overnight at room temperature or for 45 minutes at 80°C. After the APTES coating, the coverslips were rinsed twice with ethanol and twice with water. To cure the APTES layer, the coverslips were loaded in an oven for 60 min at 120 °C. The APTES coated coverslips were stored at 4 °C for a maximum of 1 week.

A solution of alginate in dH_2_O (0.5 wt%) was activated by dissolving 5 mg of 1-ethyl-3-(3-dimethylaminopropyl)carbodiimide (EDC) and 10 mg of N-hydroxysuccinimide (NHS) per ml of solution. The APTES-treated coverslips were soaked in the activated alginate for 30 minutes at RT. Excess polymer was removed and the coverslips exposed to a second aliquot of activated alginate for 90 minutes at RT, followed by rinsing three times with double-deionized water (ddH_2_O).

To functionalize the alginate surface with a DBCO linker, a 5 mM solution of DBCO-amine (Sigma Aldrich) was prepared in 20 mM sodium phosphate buffer pH 8.5 and 20% DMSO and used to dissolve 5 mg of EDC and 10 mg of NHS at no less than 5 and 10 mg per mL. A 40 µL drop of DBCO-amine was placed on parafilm and alginate surfaces were inverted on the drop and reacted for 90 minutes, followed by 3 washes with ddH_2_O.

Solutions of *peptide_S* and *peptide_I* at 200 µM were prepared in PBS and combined 1:1 (100 µM final concentration of each peptide). The peptides were placed on a shaker and allowed to hybridize for at least 1 hr prior to use. A 40 µL drop of the *peptide_S*/*peptide_I* solution was placed on parafilm, and DBCO-modified surfaces were inverted on the drop. The azide-containing *peptide_S* was reacted with the surface-bound DBCO through copper-free click chemistry for at least 2 hours. Surfaces were then washed 3 times with 1x PBS prior to use in cell adhesion or strand displacement experiments.

### Elemental composition of functionalized surfaces using X-ray photoelectron microscopy (XPS)

X-ray photoelectron microscopy (XPS) was performed on various modified surfaces to characterize each layer. Surfaces for XPS were made on silicon wafers to reduce surface charging. Wafers were prepared in the same manner as glass. Samples were run by the UNC CHANL facility. Measurements were taken on Kratos Axis Ultra DLD X-ray Photoelectron Spectrometer with monochromatic Al K α source at 150 W using a 300 x 700 μm spot size. Survey scans over a binding energy range of 0-1200 eV were taken for each sample with a constant detector pass energy range of 80 eV, followed by a high-resolution XPS measurement (pass energy 20 eV) for quantitative determination of binding energy and atomic concentration. Background subtraction, peak integration, and fitting were carried out using Kratos software. To convert peak areas to surface concentration, default instrument sensitivity factors were used (N=0.477, C=0.278, O=0.780, S=0.668, Si=0.328).

### Coiled-coil characterization using circular dichroism (CD)

Circular dichroism (CD) spectra were obtained on a Chirascan Plus Spectropolarimeter using a 10 mm path length cuvette. The peptide samples were assembled at 800 uM each, then diluted to working concentration at 65 uM in 1X PBS. CD Spectra were taken at 37 °C.

### Scanning electron microscopy (SEM) of gold nanoparticle-modified surfaces

Surfaces were prepared as described above using a biotin-modified peptide, *peptide_I-biotin*. Surfaces were incubated with gold nanoparticles modified with streptavidin (20 nm diameter, 0.5nM, Sigma) for 10 min. Excess nanoparticles were washed 3 times for 5 minutes with 1x PBS. Samples were dried and then coated with 3 nm of gold palladium alloy using a sputter coater (Cressington 108 Auto). Samples were then imaged using a FEI Helios 600 Nanolab Scanning Electron Microscope working at an accelerating voltage of 5 kV.

### Surface peptide density characterization using HRP assays

Biotin-modified surfaces were prepared using varying concentrations of *peptide_S*/*peptide_I-biotin* ranging from 0.1-100 µM. Alginate-modified surfaces were used as a no-peptide control. Surfaces prepared to test displacement were then incubated with multiple concentrations of *peptide_D* ranging from 0.1 to 60 µM in PBS, 1% DMSO for 1 h followed by 3 washes with PBS.

Following modification, surfaces were incubated with a 2 nM solution of UltraAvidin-HRP (Leinco Technologies) for 20 min at RT followed by 3x 5-min washes with 1x PBS. A solution of 2 mM ABTS and 2 mM H_2_O_2_ was added to the coverslips and allowed to react for 6 hrs in the dark. Following this, absorbance was measured at 450 nm using a Thermo Fisher Varioskan Lux Plate Reader and compared to standards to determine the concentration of peptides bound to the surface.

### Scanning electron microscopy (SEM) of cells on displayed and displaced surfaces

For SEM experiments, cells were cultured and manipulated as described above to illustrate different conditions/surfaces. Cells were then fixed with 2% PFA, 2.5% glutaraldehyde in PBS for 1 h at room temperature. Fixed samples were dehydrated by exposure to a graded series of water-ethanol mixtures. Once in 100% ethanol, samples were dried at the critical point of CO_2_ using a critical point drying technique (Tousimis Samdri-795) to preserve structural details. Dried samples were then coated with 10 nm of gold palladium alloy using a sputter coater (Cressington 108 Auto), and imaged using a FEI Helios 600 NanolabScanning Electron Microscope working at an accelerating voltage of 5 kV.

### Fibroblast adhesion assays

NIH 3T3 mouse embryonic fibroblasts (ATCC CRL-1658) were cultured in DMEM/F12 (1:1) media (Gibco) with 10% FBS (VWR), 1% antibiotic/antifungal solution (Gibco) and passaged every 3 days. HEK-293 cells (Human embryonic kidney, ATCC-CRL-1573™), used as a control cell line, were cultured in DMEM, high glucose (Gibco) supplemented with 10% FBS and 1% AA.

For adhesion experiments, 3500-5000 cells were plated on functionalized surfaces in 24-well plates and allowed to adhere and spread for the desired time (typically 2-4 hours) at 37 °C and 5% CO2. Adhesion assays were carried out under reduced-serum conditions (2% FBS). Following this incubation, cells were fixed for 15 min in 4% PFA or displaced by the addition of 60 µM *peptide_D* for the desired time (typically 1 hr) and then fixed with 4% PFA in PBS for 15 min. Cells were permeabilized in 0.2% Triton-100 for 5 min. Following permeabilization, cells were washed three times with cold PBS containing 1 mM CaCl_2_ and then incubated with Alexa-fluor phalloidin (Thermo) at 1:200 in PBS for 1 hr at RT or overnight at 4 °C. Following this, cells were washed 3 times with PBS and incubated with DAPI (Sigma) at 1:1000 in PBS for 20 min. Cells were then washed 3 times in PBS and mounted on glass slides using Thermo Immu-MountImmumount. Coverslips were imaged on the GE INCell Analyzer 2200 high-content microscope.

### Immunocytochemistry and confocal fluorescence microscopy

The following primary antibodies were used for immunocytochemistry; mouse anti-Vinculin (1:500, Sigma Aldrich V9131), rabbit anti-Paxillin [Y113] (1:500, Abcam ab32084), rabbit anti-β_3_/CD61 [SJ19-09] (1:500, Invitrogen MA5-32077), and rat anti-β^1^ [MB1.2] (1:250, Sigma MAB1997).

For confocal microscopy, cells were prepared as described above. Following fixation and permeabilization, coverslips were washed three times with PBS and then blocked for 30 minutes with 1% Bovine Serum Albumin (BSA, Sigma) in PBS + 0.2% Tween20 (PBS-T). After blocking, coverslips were incubated with primary antibodies diluted in PBS-T with 1% BSA for 1hr at RT or overnight at 4°C. Coverslips were washed with PBS-T three times for 5 minutes followed by the addition of the appropriate Alexa Fluor 488, 568, and/or 647 conjugated secondary antibodies (1:1000, Invitrogen) or Alexa Fluor 647-Phalloidin (1:400, Invitrogen) diluted in PBS-T with 1% BSA 1 h at RT or overnight at 4°C. Following this, cells were washed 3 times in PBS-T then incubated with DAPI (Sigma) 1:1000 or Hoechst (Sigma) 1:2000 in PBS for 20 min. Cells were then washed and mounted on glass slides using Thermo Immu-Mount and imaged on a Ziess LSM 880 Confocal Microscope.

### Integrin binding ELISA

#### Plate-Modification

Flat-bottom 96-well ELISA plates (Nunc, MaxiSorp) were coated with 1.0 µg/mL integrin (R&D, human α_V_β_3_or α_5_β_1_, 100µL/well) in 50 mM carbonate buffer pH 9.6 for 2 hours at 37°C. Plates were washed 3X with PBS-T (1X PBS, 0.01% Tween 20; 5min, 200µL/well, room temperature). Plates were blocked twice with TS-B (20 mM Tris-HCl pH 7.5, 150 mM NaCl, 1mM CaCl2, 1 mM MgCl2, 1 mM MnCl2, 1% BSA; 1 hour, 200µL/well, room temperature), and washed 3X with PBS-T.

## Competitive Binding Assays

*Peptide_S* and *peptide_I* were pre-hybridized at a 1:1 ratio in TS-B for 2 hours at room temperature. Human fibronectin (Sigma-Aldrich, 1.0 µg/mL in TS-B) was added to integrin-coated plates with serial dilutions of peptide (10 µM to 3.2 nM) and allowed to bind for 30 minutes at room temperature (100 μL/well). The plate was washed 3x with PBS-T (5 min each, 200uL/well) to remove unbound fibronectin and peptide. Primary antibody (Santa Cruz Biotechnology, mouse monoclonal anti-fibronectin, 1.0 µg/mL in TS-B) was added to the wells for 1 hour at room temperature. The plate was then washed 3x with PBS-T (5 min each, 200μL/well) to remove unbound antibodies. A secondary HRP-conjugated antibody (Invitrogen, goat anti-mouse IgG(H+L) HRP, 2.0 µg/mL) was added for 1 hour at room temperature. All wells were washed 3x with PBS-T (5 min, 200uL/well) and HRP reactions were initiated by the addition of 50 μL TMB substrate solution (RayBiotech). Color was developed for 30 minutes before reactions were quenched by the addition of 50 µL TMB Stop Solution (RayBiotech). Absorbance was measured at 450 nm using a Thermo Fisher Varioskan Lux Plate Reader. IC50 values were determined using GraphPad Prism 9.0.0 by fitting the data to the Hill Equation.

## Cell adhesion kinetics and dosing experiments

Cells were plated on modified coverslips as described above. All samples were run in duplicate. Cells were allowed to adhere for 0.5, 1, 2, or 3 h on RGDS surfaces. Cells from each timepoint were washed once with PBS and fixed with 4% PFA in PBS, and additional coverslips were used for the displacement experiments. For displacement, after cells adhered for the desired time, 300 µL of culture media was removed and *peptide_D* was added to a final concentration of 60 µM, 1% DMSO. Cells were displaced for 1 hour, followed by one wash in PBS and fixation in 4% PFA. Two additional coverslips were displaced using 1 µM *peptide_D* for 1 hour. After displacement, surfaces were washed 1x with PBS, followed by the addition of 100 µM *peptide_I* in cell culture media. Cells were allowed to re-adhere for 2 h and then fixed with 4% PFA in PBS.

### Live cell imaging and ON-OFF signaling over multiple cycles

For live cell experiments, 35 mm glass bottom dishes (Mattek) were modified instead of coverslips. Cells were plated on the modified dishes and appropriate spots were located using the built-in Metamorph software. Cells were allowed to spread for the desired time, followed by the addition of 60 µM *peptide_D* for 45 min-1 hr. Half the media was removed and then *peptide_I* was added to a final 100 µM. Cells were allowed to re-adhere for the desired time and then the process was repeated. Live cell imaging was performed using an Olympus VivaView long-term live-cell wide field microscope at 37°C, 5% CO_2_. Images were taken every 15-30 seconds.

### Cell viability over multiple ON-FF cycles

Assessment of cell viability was performed with PBS containing 2 μM calcein-AM (Life Technologies) and 4 µM propidium iodide (Sigma) for 20 min at 37 °C. The cells were then rinsed with PBS and imaged using INCELL Analyzer 2200 (GE Technologies). Cells were then manually counted in the red and green channels and percent viability was determined from the ratio of green/total cell counts.

### Data Analysis and Statistics

#### Cell size quantification on stained cells

Fluorescent images were loaded into Fiji and threshold using standard procedures. The “analyze particles” function was then used to measure cell surface area Cell number was quantified through counting nuclei using DAPI staining.

For quantification of fluorescently stained samples, a minimum of 500 randomly selected cells from a minimum of three independent batches of culture were included for each condition. Morphometric, quantitative, live-image and displacement analysis of images and all related quantifications were performed using ImageJ software (National Institute of Health). Cell counts and cell surface area were determined on threshold images with adjustments for contrast, brightness, and color balance to obtain optimum visual reproduction of data.

### DIC image segmentation

DIC images were first corrected for illumination gradients. An illumination map of the raw image was calculated in MATLAB by implementing a 10-step Alternating Sequential Filter.^33^ This illumination map was then applied to the original raw image using Cell Profiler. This result was used for cell segmentation.

Segmentation of illumination-corrected DIC images was performed using the u-net Segmentation plugin on imageJ.^34^ A new segmentation model was trained for more than 10,000 iterations using 10 manually segmented representative images from the collected videos. Following this, the segmentation model was applied to each video and the cell masks calculated. Frames where the segmentation was obviously incorrect were not used in the following calculations of cell area and spreading kinetics.

### Cell spreading rate analysis

The rate of change in cell area was determined by calculating the slope over the first 6 minutes after each cell began to respond to the added peptide (*peptide_I* or *peptide_D*). Cells that did not respond to peptide addition were removed from the analysis.

### Focal Adhesion and Enrichment Ratio

Methods adapted from Spatz et al.^35^ Immunofluorescence images were first blurred using a gaussian threshold (diameter 2 pixels). The cell border was determined through applying an automatic threshold (Huang method) of the Vinculin channel. Focal adhesions were then identified through an automatic threshold using a Max Entropy algorithm. To remove individual bright pixels, a threshold of 0.1 square microns was used. The mean grey value of the focal adhesions was then measured and divided by the mean grey value of the remaining region of interest. For analysis of size, number, and aspect ratio of focal adhesions, a minimum filter (diameter 5 pixels) was used to reduce noise in the center of the cell before using an automatic threshold determined by the Otsu method.

### Statistical analysis

Statistical analysis was performed using Graphpad Prism 9 software. For the determination of statistical significance, a student’s t-test or Kruskall-Wallis test by ranks was implemented. P values<0.05 were deemed significant. Values in graphs are shown as mean±s.d (standard deviation). Comparison between multiple groups was done by implementing two-way Analysis of variance (ANOVA) with a Bonferroni coefficient post hoc correction.

## Associated content

### Notes

The authors declare no competing financial interest.

## Acknowledgements

We thank the Department of Applied Physical Sciences at UNC-Chapel Hill for support of this work. M.L.D. acknowledges support from the NSF Graduate Research Fellowship Program. Mass Spectrometry was performed at the UNC Mass Spectrometry Core Laboratory at CRITCL (Chemical Research Instrumentation Teaching and Core Labs) in the Department of Chemistry. XPS and SEM work was performed at the Chapel Hill Analytical and Nanofabrication Laboratory (CHANL), a member of the North Carolina Research Triangle Nanotechnology Network (RTNN) which is funded by the National Science Foundation (NSF) (Grant ECCS-1542015) as part of the National Nanotechnology Coordinated Infrastructure (NNCI). CD measurements were taken at the UNC Macromolecular Interactions Facility of the Lineberger Comprehensive Cancer Center, funded by the National Cancer Institute of the National Institutes of Health (NIH) (P30CA016086). Confocal Microscopy was performed at the UNC Hooker Imaging Core Facility, supported in part by P30CA016086 Cancer Center Core Support Grant to the UNC Lineberger Comprehensive Cancer Center. We thank the Be A Maker (BeAM) network of makerspaces at UNC Chapel Hill for the use of the laser cutter.

## Supplementary Information

## Supplementary Peptide Characterization, Schematics and Experimental Data

Table S1

S1- Peptide_I

S2- Peptide_S

S3- Surface synthesis scheme

S4- Peptide_D

S5- XPS atomic composition

S6- CD of Peptide_D

S7- Peptide_I-biotin

S8- SEM of cells on displayed and displaced surfaces

S9- Peptide_I-RGES

S10- ELISA

S11- Focal adhesion/integrin staining of negative controls

S12- Cell area with time of display

S13- Explanation of ER calculation

S14- Kinetics of cell spreading on APTES, Glass, and RGDS

S15- CD kinetics of displacement in solution

S16- Cell number over ON-OFF cycles

S17- Cell viability over ON-OFF cycles

## Videos

S1- Cells spreading on RGDS

S2- Cells on alginate

S3- Cells on Peptide_S

S4- HEK cells on RGDS

S5- Cell spread and contract over multiple cycles

S6- Cells on glass

S7- Cells on APTES

## Peptide Characterization

**Table S1.**
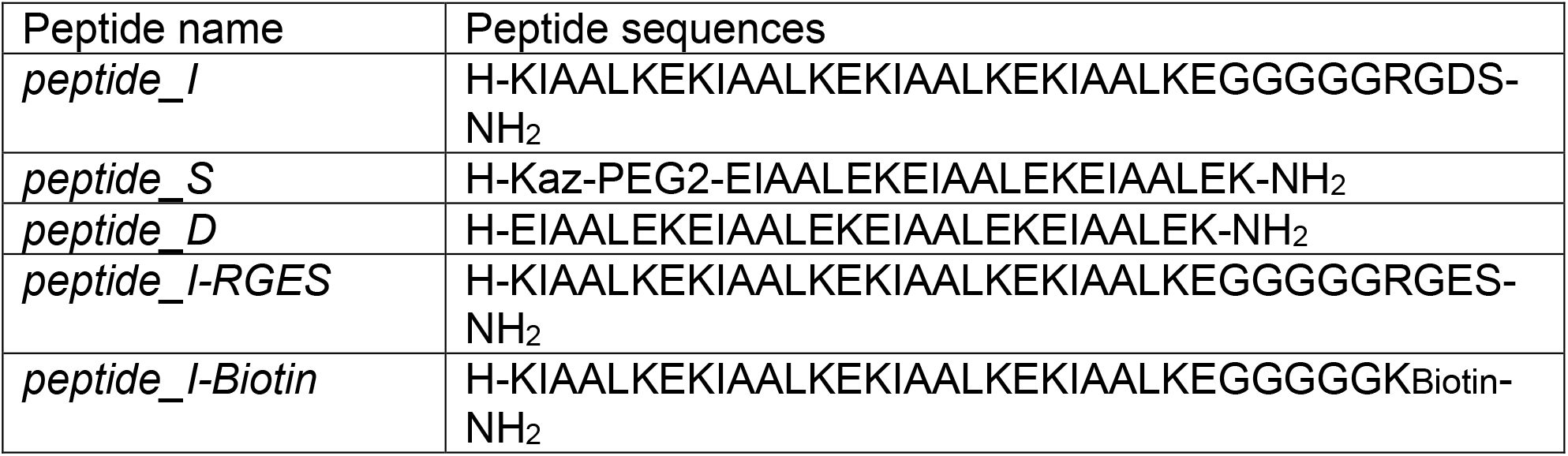
Sequences of all peptides used. Peptide names and corresponding peptide sequences used.

| Peptide name | Peptide sequences |
| --- | --- |
| *peptide_I* | H-KIAALKEKIAALKEKIAALKEKIAALKEGGGGGGRGDS-NH<sub>2</sub> |
| *peptide_S* | H-Kaz-PEG2-EIAALEKEIAALEKEIAALEK-NH<sub>2</sub> |
| *peptide_D* | H-EIAALEKEIAALEKEIAALEKEIAALEK-NH<sub>2</sub> |
| *peptide_I-RGES* | H-KIAALKEKIAALKEKIAALKEKIAALKEGGGGGGRGES-NH<sub>2</sub> |
| *peptide_I-Biotin* | H-KIAALKEKIAALKEKIAALKEKIAALKEGGGGGK Biotin-NH<sub>2</sub> |

**Figure S1.**
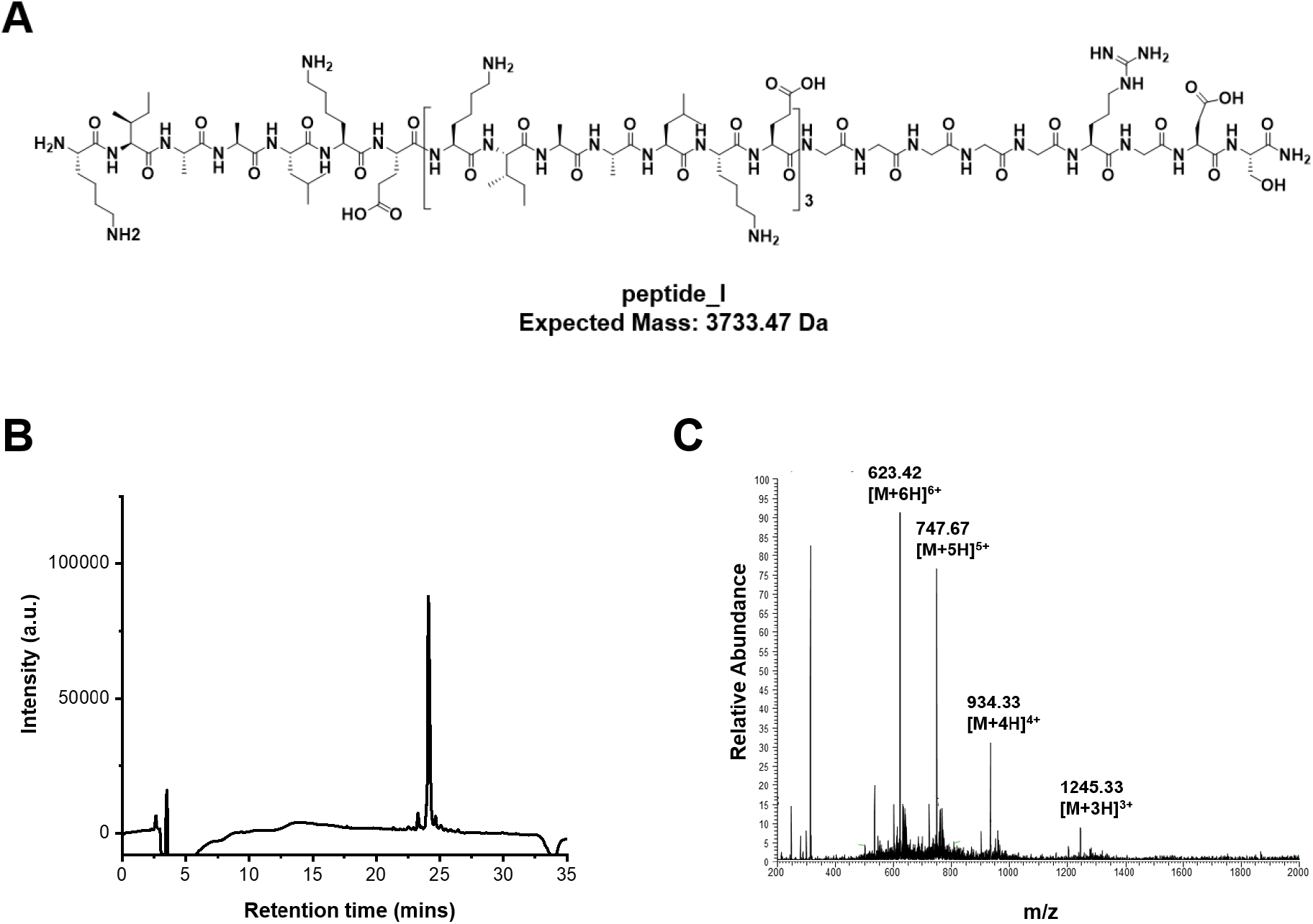
Structure and HPLC-MS characterization of peptide_I. (A) Chemical structure and expected mass of *peptide_I.* (B) Analytical HPLC (monitoring peptide absorbance at 214 nm) and (C) ESI-MS confirmed the peptide identity and high purity.

**Figure S2.**
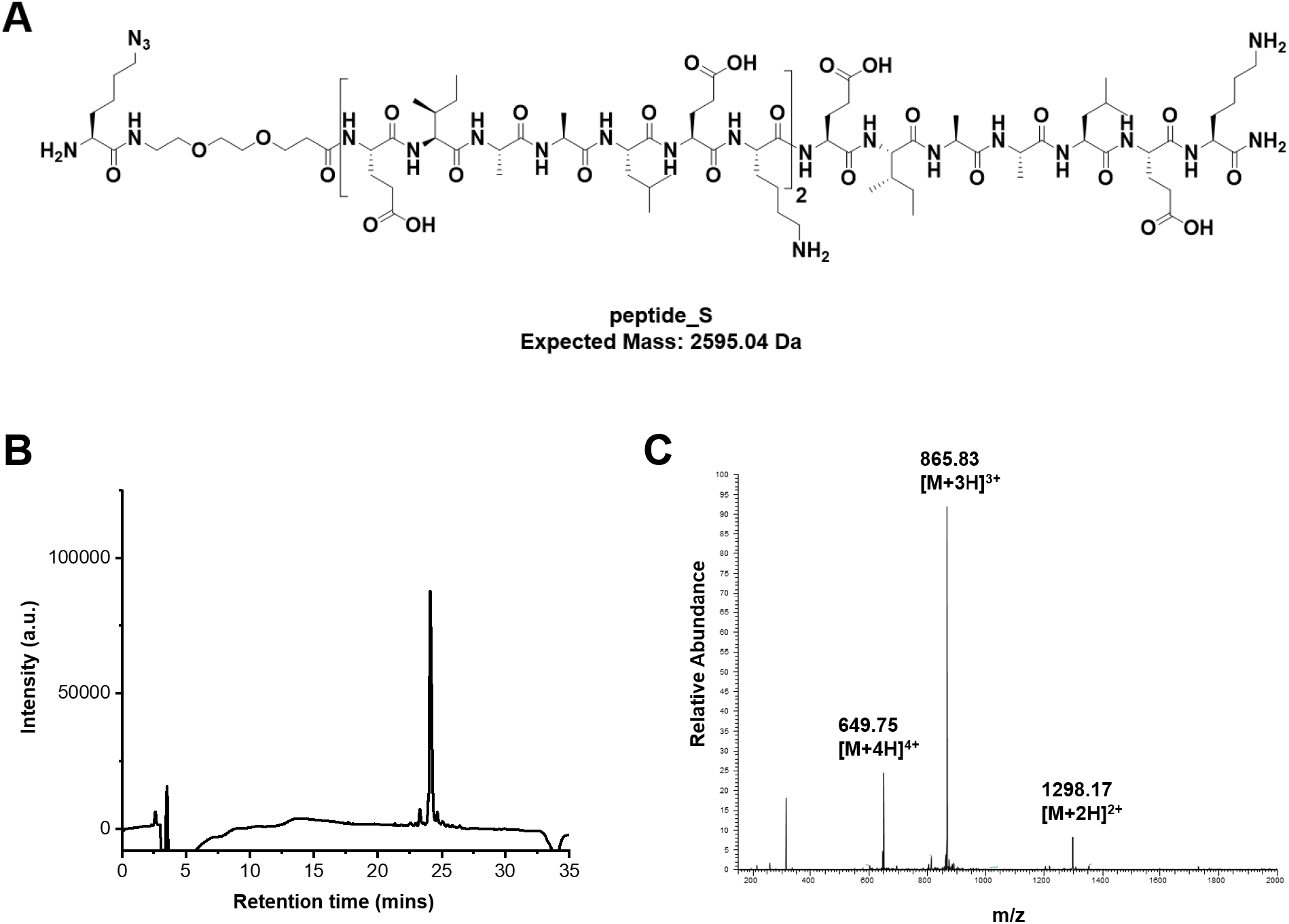
Structure and HPLC-MS characterization of peptide_S. (A) Chemical structure and expected mass of *peptide_S* (K^az^-PEG_2_- EIAALEKEIAALEKEIAALEK). (B) Analytical HPLC (monitoring peptide absorbance at 214 nm) and (C) ESI-MS confirmed the peptide identity and high purity.

**Figure S3.**
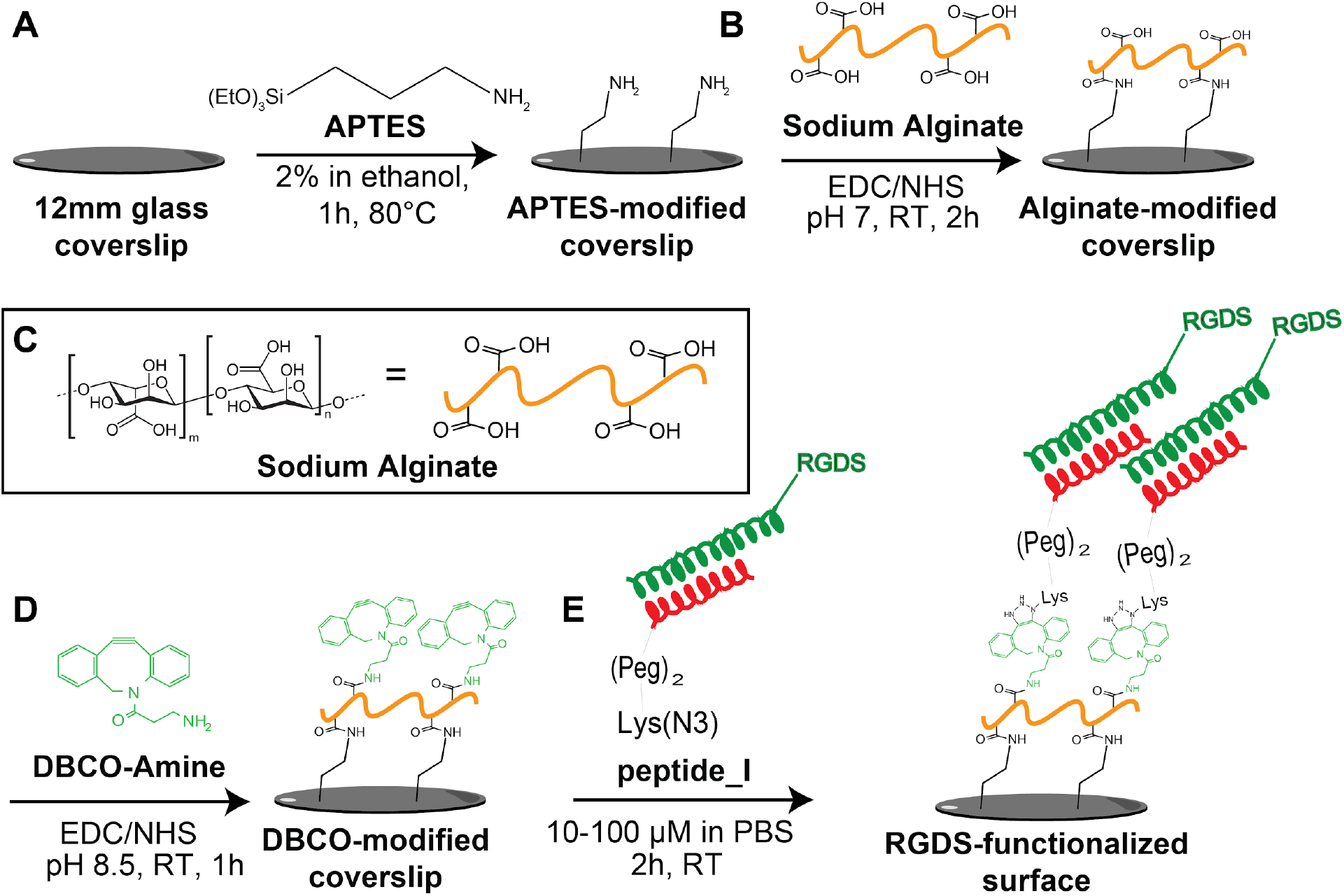
Schematics of surface modification method. (A) Glass coverslips were coated with (3-Aminopropyl)triethoxysilane (APTES). (B) APTES surfaces were modified with a layer of sodium alginate (0.5 wt%) through an EDC/NHS conjugation. (C) Chemical formula of sodium alginate. (D) Alginate surface was modified with amino-DBCP through an EDC/NHS reaction (E) *peptide_S*/*peptide_I-RGDS* hybrid was added to the DBCO surface using copper-free click chemistry.

**Figure S4.**
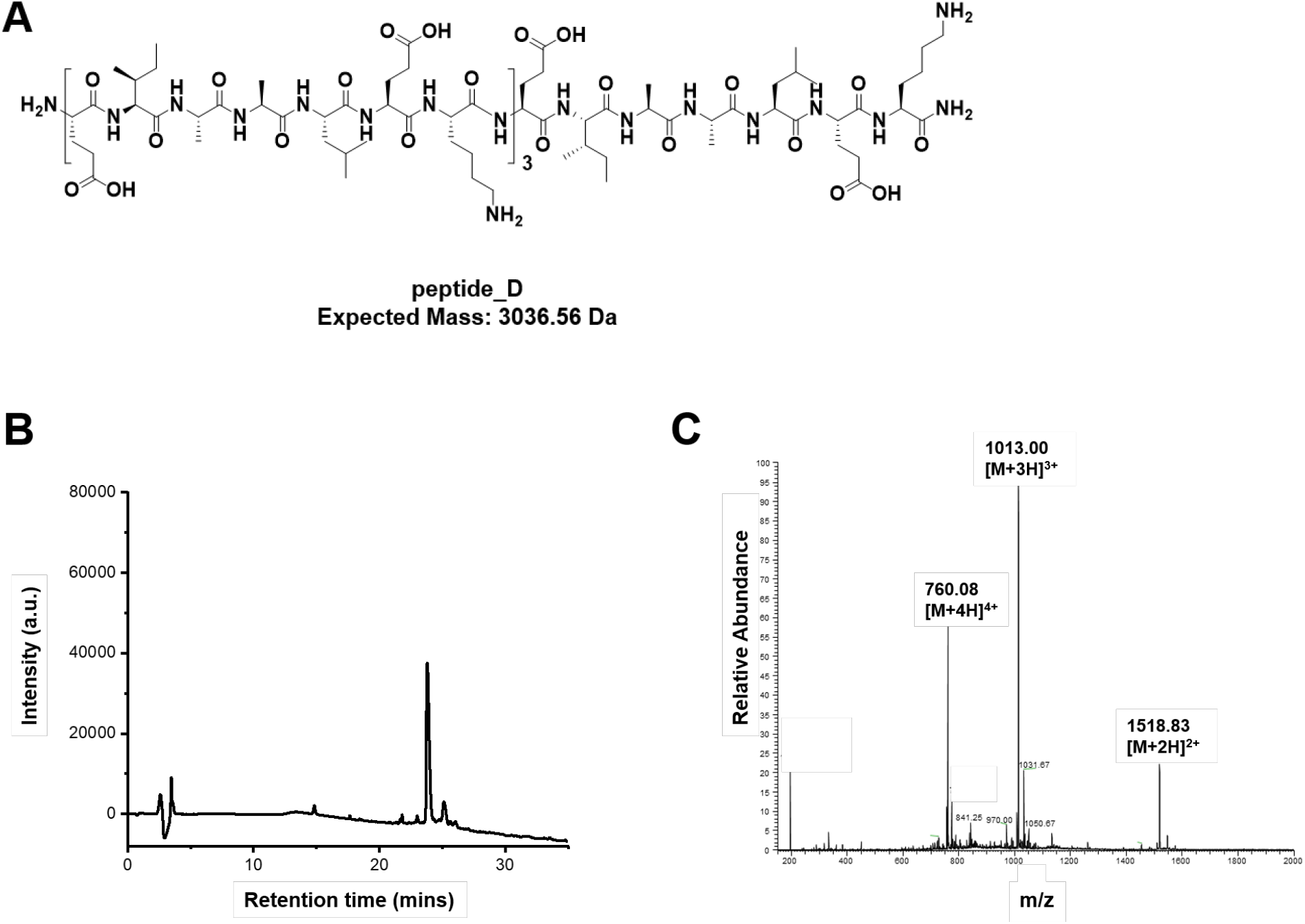
Structure and HPLC-MS characterization of peptide_D. (A) Chemical structure and expected mass of *peptide_D* (EIAALEKEIAALEKEIAALEKEIAALEK). (B) Analytical HPLC (monitoring peptide absorbance at 214 nm) and (C) ESI-MS confirmed the peptide identity and high purity.

**Figure S5.**
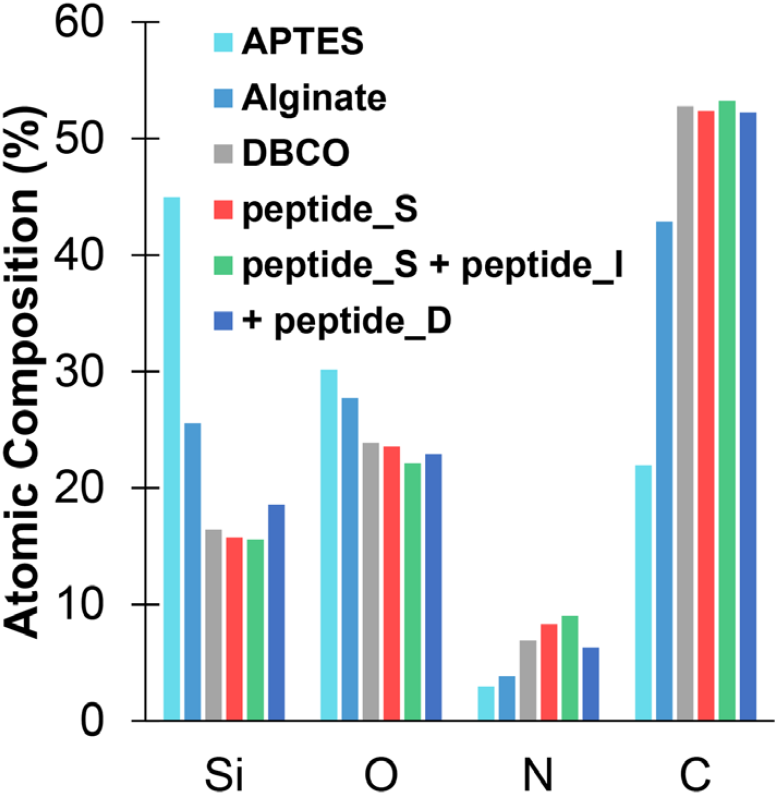
XPS characterization of the atomic composition of the modified surfaces. Percent atomic composition of the surface from XPS data obtained at each layer of chemical modification. Values were used to compute the C/O and C/N ratios described in **Figure 1C**. A decrease in percent Si and O suggests layering on the glass substrate. A progressive increase in percent N and C from APTES to alginate to DBCO to *peptide_S* suggests successful chemical layering at each step to modify the surface with peptide.

**Figure S6.**
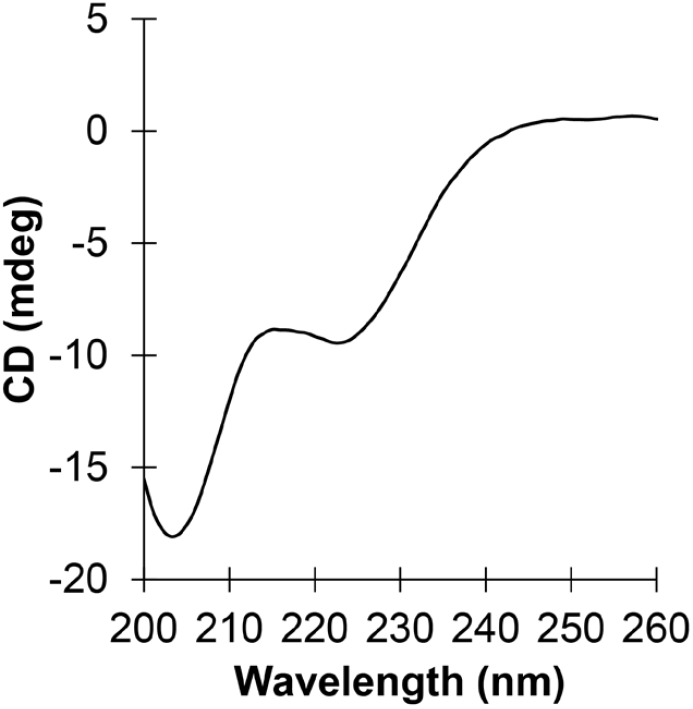
Circular dichroism of peptide_D. The spectrum is typical for α-helix peptides; the minima at 222 nm and a larger minima at 208 nm suggest that the peptide is forming an α-helix.

**Figure S7.**
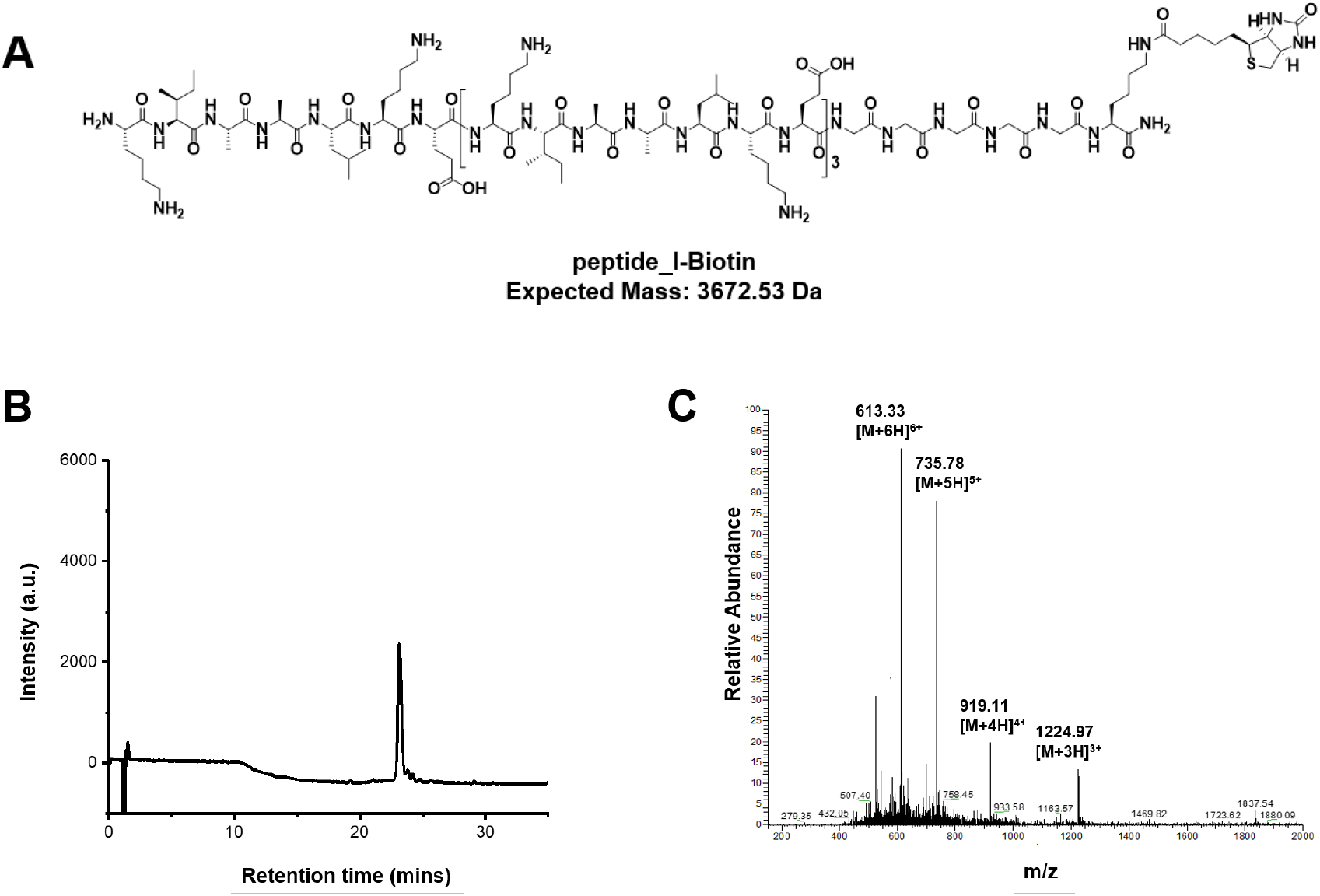
Structure and HPLC-MS characterization of peptide_I-biotin. (A) Chemical structure and expected mass of *peptide_I-biotin* (KIAALKEKIAALKEKIAALKEKIAALKEGGGGGK(biotin)). (B) Analytical HPLC (monitoring peptide absorbance at 214 nm) and (C) ESI-MS confirmed the peptide identity and high purity.

**Figure S8.**
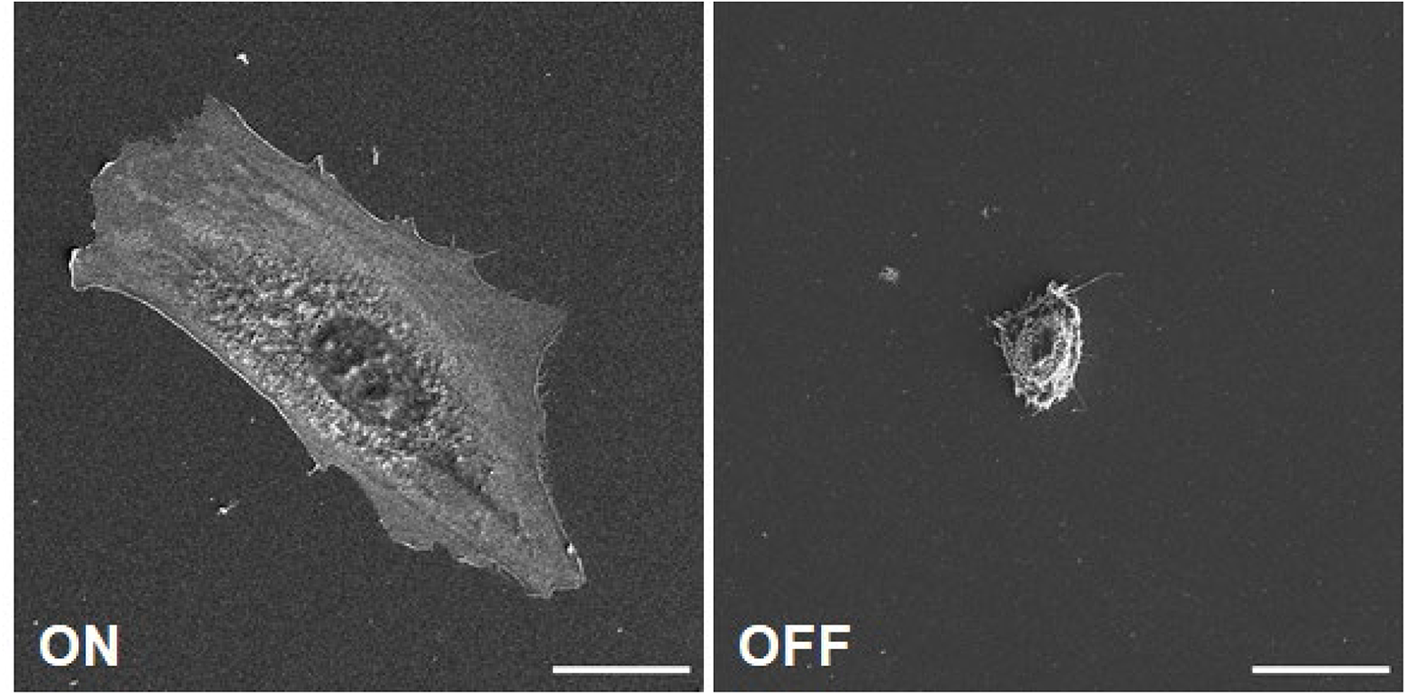
Scanning electron microscopy images of a 3T3 cell cultured on RGDS-displayed (left) and displaced (right) surfaces (scale bar: 20 μm).

**Figure S9.**
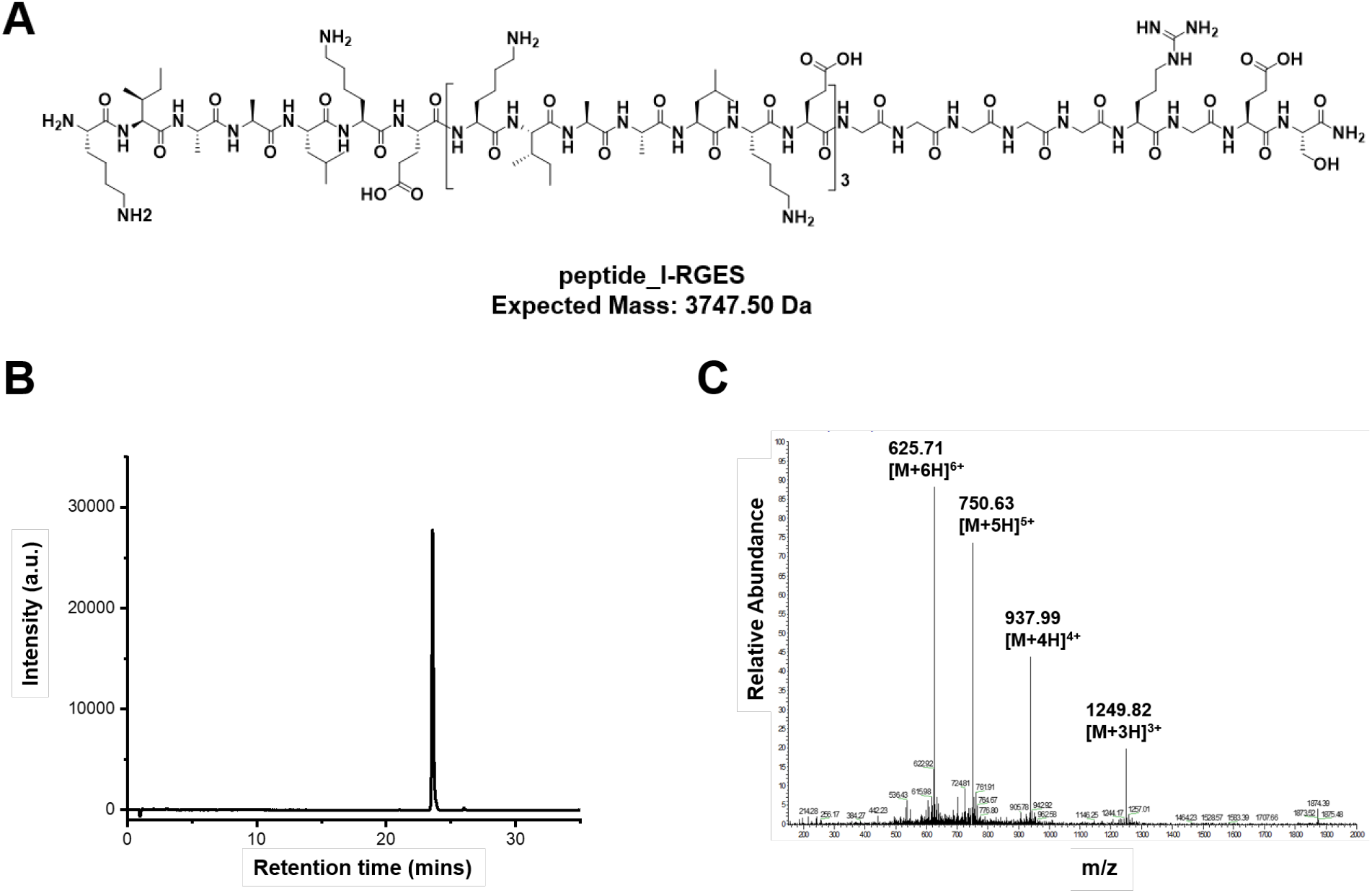
Structure and HPLC-MS characterization of peptide_I-RGES. (A) Chemical structure and expected mass of *peptide_I-RGES* (KIAALKEKIAALKEKIAALKEKIAALKEGGGGGRGES). (B) Analytical HPLC (monitoring peptide absorbance at 214 nm) and (C) ESI-MS confirmed the peptide identity and high purity.

**Figure S10.**
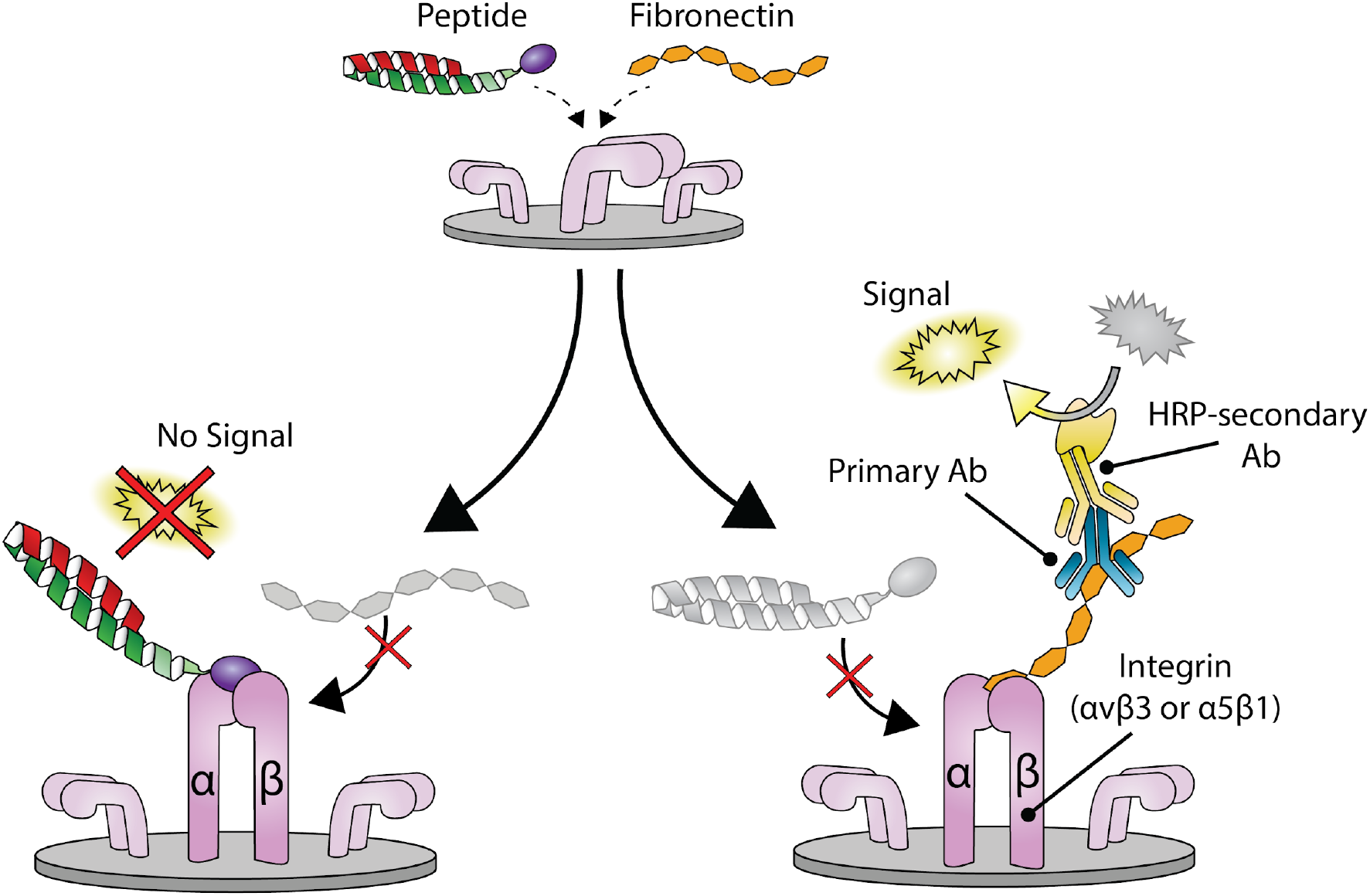
Schematic of the ELISA experiments. Integrin modified plates are incubated with fibronectin and varying concentrations of RGDS peptide. When the peptide inhibits fibronectin binding to the integrins (left), no signal is detected. Without peptide inhibition (right), fibronectin binds the integrins and is then detected using HRP-modified antibodies.

**Figure S11.**
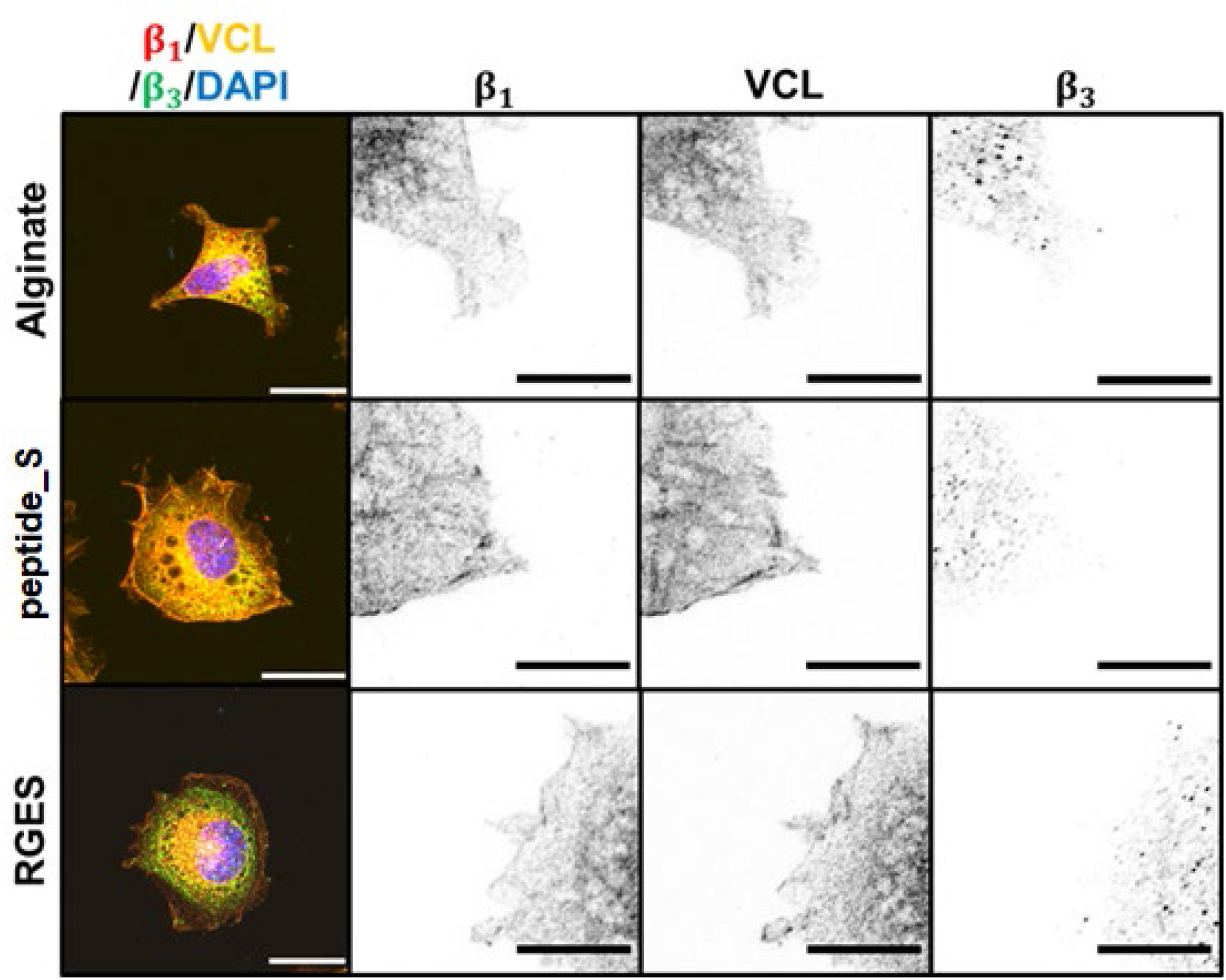
Cell adhesion on control surfaces. Cells cultured on surfaces of Alginate, *peptide_S*, and RGES did not display significant focal adhesions or β1/β3 colocalization

**Figure S12.**
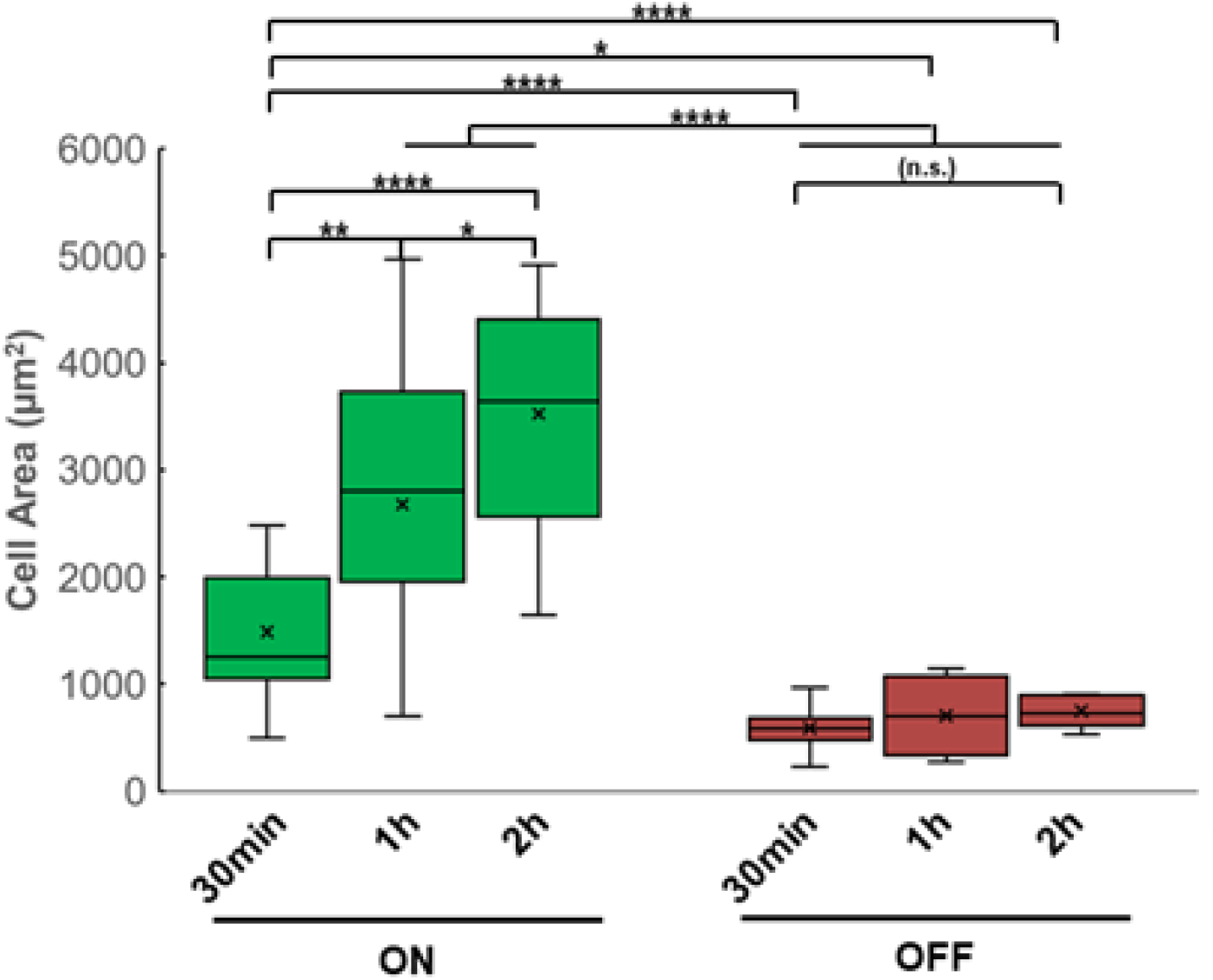
Cell area as a function of signal display time before displacement. P-Values: *<0.05, **<0.01, ***<0.001, ****<0.0001.

**Figure S13:**
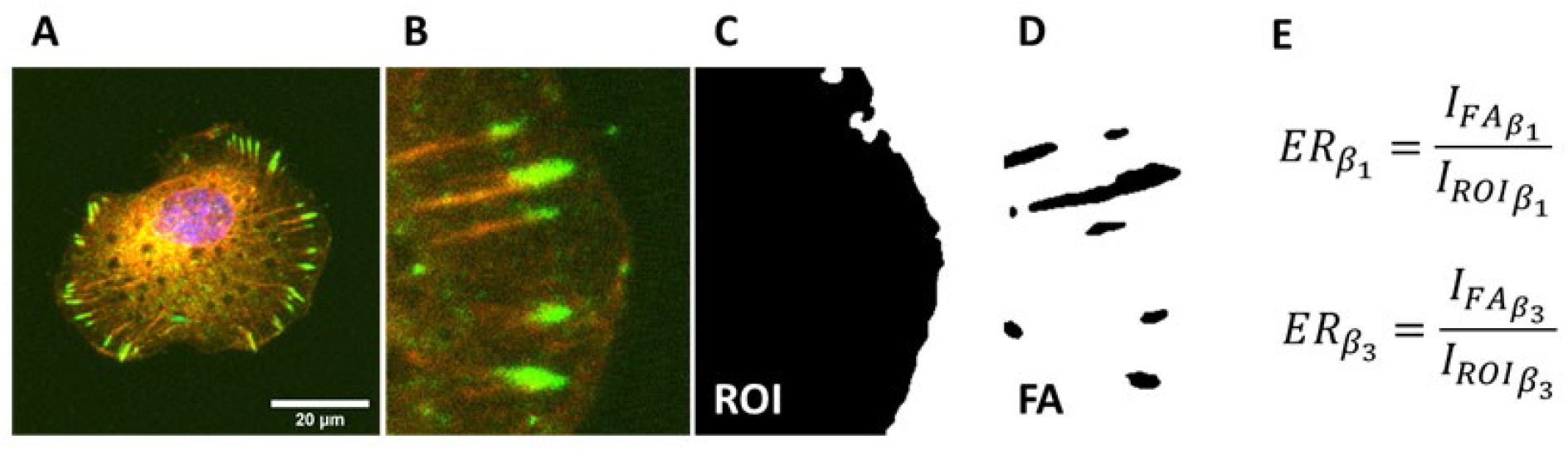
Analysis of integrin enrichment ratio. (A) CLSM image of cell and region of interest selected. (C) Area of the cell determined automatically and (D) area of the focal adhesions determined automatically through vinculin staining from the region of interest. (E) Measurements and calculation of enrichment ratio. Intensity of focal adhesions was not included in the ROI measurement.

**Figure S14.**
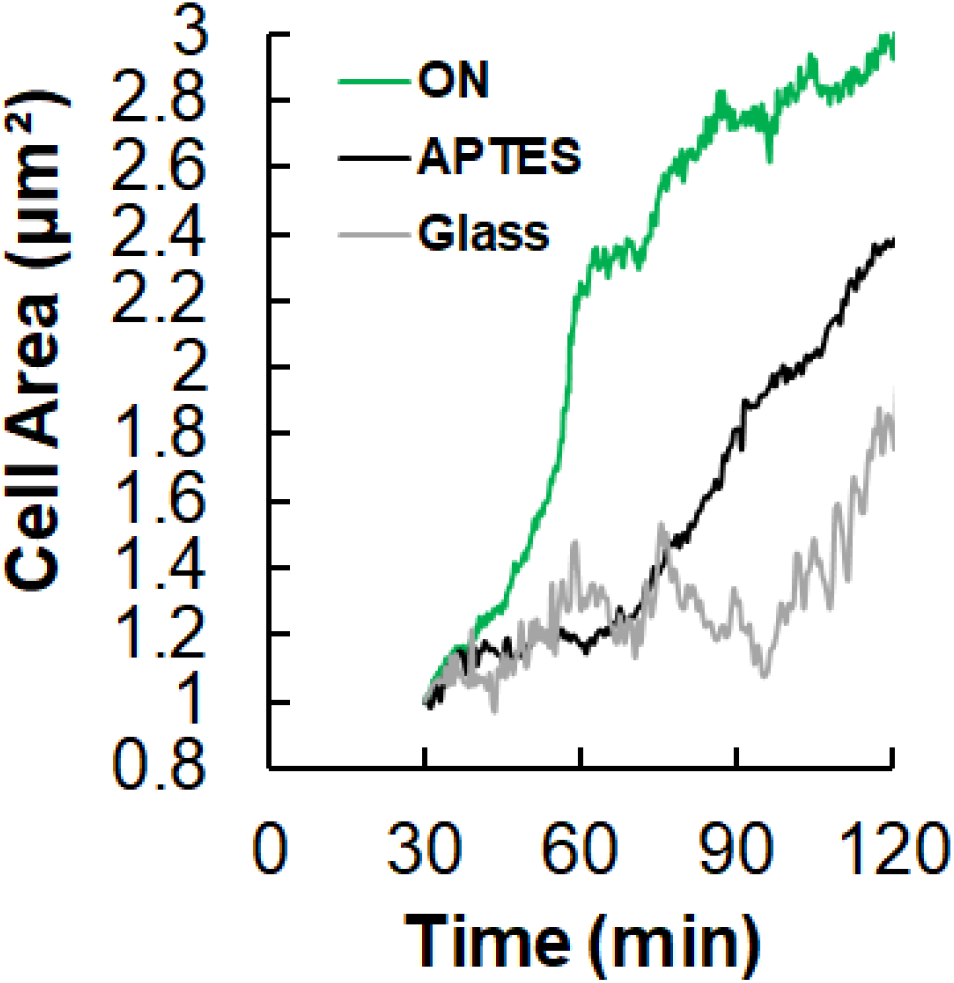
Cell spreading kinetics on surfaces displaying peptide_I compared to controls. Increase in average cell area over time on a surface in the ON state compared to an APTES-coated surface and glass. Kinetic curves of cell spreading were normalized to the initial cell area.

**Figure S15.**
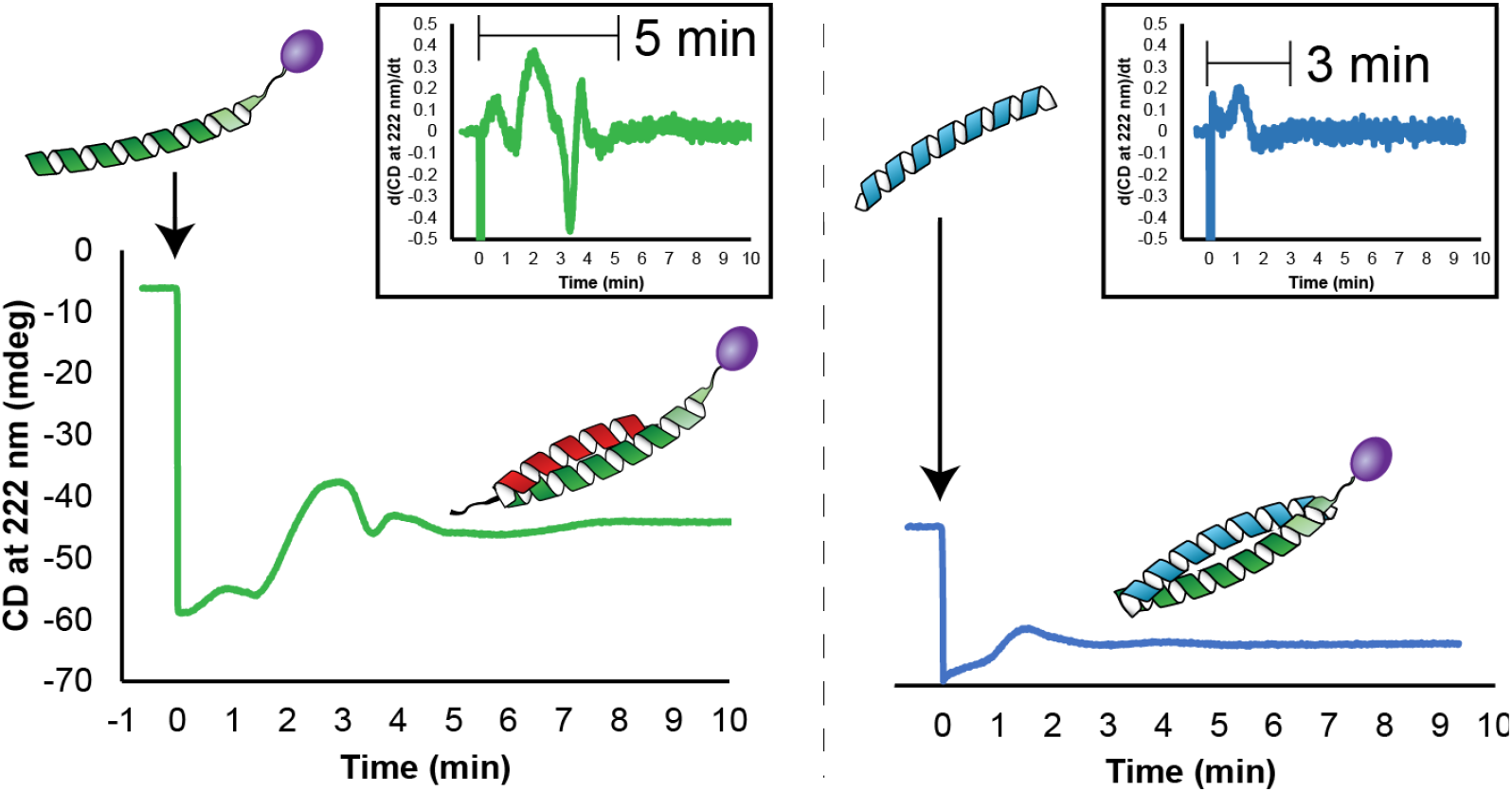
Coiled-coil behavior of signal addition and displacement. Circular dichroism of peptides in solution, diluted in PBS to 60 μM at 37 °C to mimic cell testing conditions. CD signal at 222 nm over time of *peptide_S* with subsequent additions of *peptide_I* (green) followed by the displacer *peptide_D* (blue). Insets show the derivative of the CD signal at 222 nm over time, showing the time it takes to form the *peptide_S*/*peptide_I* dimer (green; 5 min) and the time it takes to displace *peptide_S* and form the *peptide_I*/*peptide_D* dimer (blue; 3 min).

**Figure S16.**
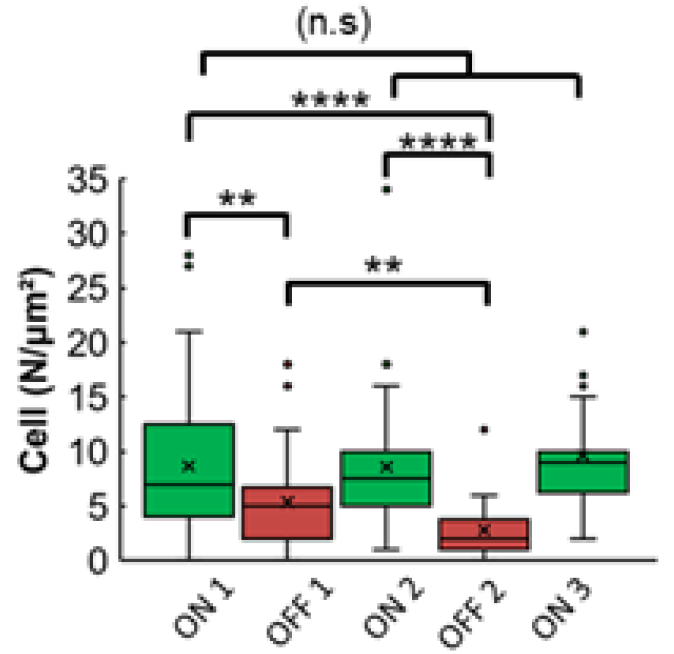
Number of cells per micrometer squared at each cycle step. P-Values: *<0.05, **<0.01, ***<0.001, ****<0.0001.

**Figure S17.**
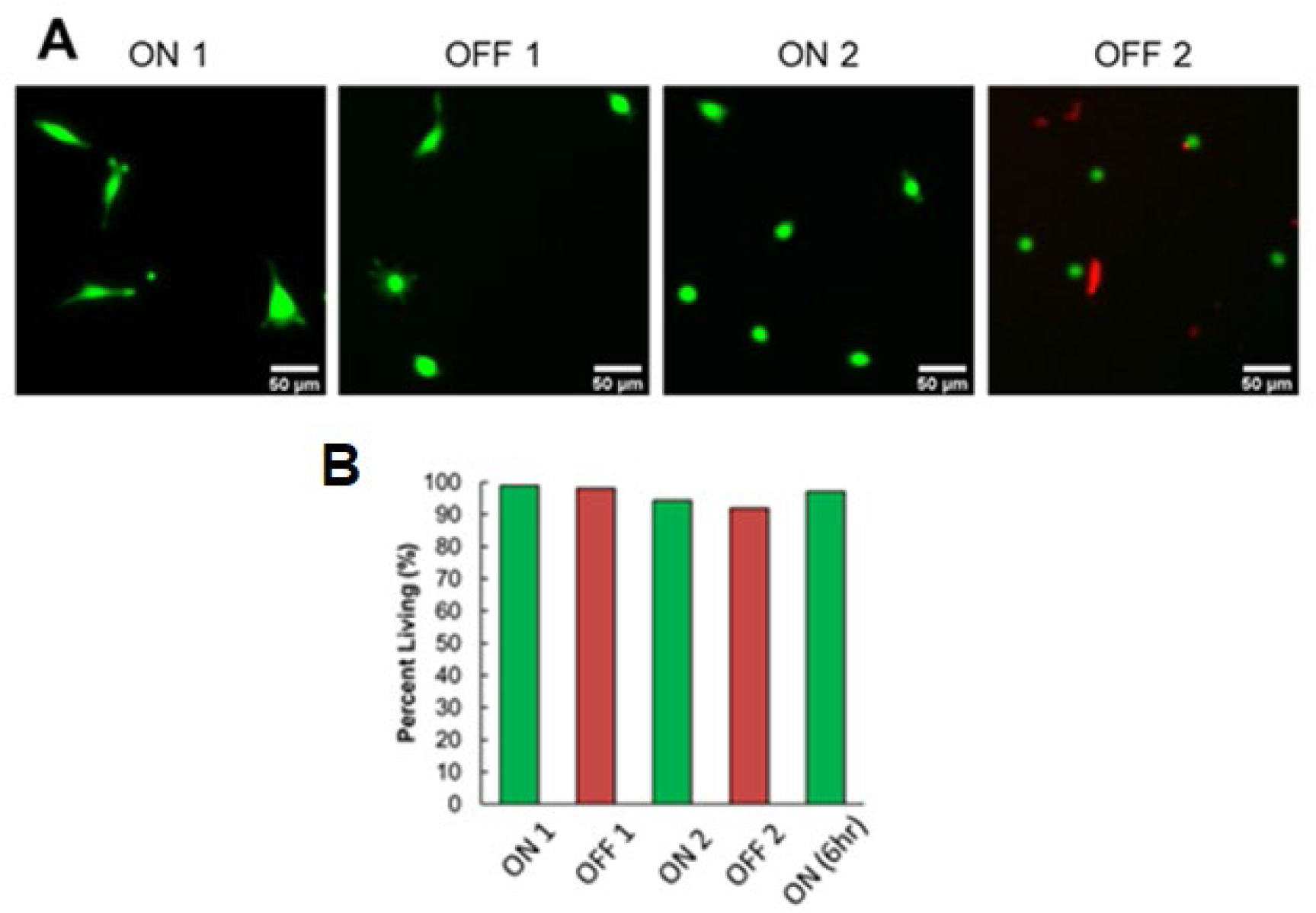
Live-dead assay to determine cell viability. (A) Representative confocal microscopy images. (B) Quantification of cell viability from confocal microscopy shows >90% cell viability with all display-displace cycles.

## Supplementary Videos

**Video S1**. *Video of NIH 3T3 cells spreading on a peptide_I-RGDS surface* 20x DIC video of NIH 3T3 cells plated on fully modified peptide_I-RGDS surface. Time stamp represents time since the video started. Cells were plated 30 minutes prior to the start of the video; images taken every 30sec.

**Video S2**. *Video of NIH 3T3 cells on an alginate surface*

10x DIC video of NIH 3T3 cells plated on a surface modified only with alginate. The non-fouling surface shows little to know spreading without bioactive RGDS signal present. Time stamp represents time since the video started. Cells were plated 30 minutes prior to the start of the video; images taken every minute.

**Video S3**. *Video of NIH 3T3 cells on a surface modified with peptide_S*

10x DIC video of NIH 3T3 cells plated on a surface modified only with peptide_S. The negative control surface shows little to know spreading without bioactive RGDS signal present. Time stamp represents time since the video started. Cells were plated 30 minutes prior to the start of the video, images taken every minute.

**Video S4**. *Video of HEK 293T cells on a peptide_I-RGDS surface*

10x DIC Video of HEK 293T cells plated on a peptide_I-RGDS modified surface. Cells are significantly less responsive to the RGDS signal due to lack of integrin α_5_β_1_ expression and show little to know adhesion on the surface. Time stamp represents time since the video started. Cells were plated 30 minutes prior to the start of the video; images taken every minute.

**Video S5**. *Video of cells in response to signal display and displacement*

10x DIC Video of NIH 3T3 cells plated on a peptide_I-RGDS modified surface. Cells spread and contract with addition of peptide signals (peptide_I-RGDS for “ON” and peptide_D for “OFF”). Timepoints of signal addition and displacement are labeled on the video. Images taken every 15 seconds; time stamp shows time since plating. Scale bar = 100μm

**Video S6**. *Video of NIH 3T3 cells spreading on glass*

20x DIC video of NIH 3T3 cells plated on a bare, plasma-etched glass coverslip (negatively-charged). Time stamp represents time since the video started. Cells were plated 30 minutes prior to the start of the video; images taken every 30sec.

**Video S7**. *Video of NIH 3T3 cells spreading on APTES*

20x DIC video of NIH 3T3 cells plated on an APTES coated surface (positively-charged). Time stamp represents time since the video started. Cells were plated 30 minutes prior to the start of the video; images taken every 30sec.

## References

(1) Frantz, C., Stewart, K. M., Weaver, V. M. The Extracellular Matrix at a Glance. J. Cell Sci. 2010, 123 (24), 4195–4200. https://doi.org/10.1242/jcs.023820.

(2) Theocharis, A. D., Skandalis, S. S., Gialeli, C., Karamanos, N. K. Extracellular Matrix Structure. Adv. Drug Deliv. Rev. 2016, 97, 4–27. https://doi.org/10.1016/j.addr.2015.11.001.

(3) Singh, P., Carraher, C., Schwarzbauer, J. E. Assembly of Fibronectin Extracellular Matrix. Annu. Rev. Cell Dev. Biol. 2010, 26 (1), 397–419. https://doi.org/10.1146/annurev-cellbio-100109-104020.

(4) Midwood, K. S., Williams, L. V., Schwarzbauer, J. E. Tissue Repair and the Dynamics of the Extracellular Matrix. Int. J. Biochem. Cell Biol. 2004, 36 (6), 1031–1037. https://doi.org/10.1016/j.biocel.2003.12.003.

(5) Madl, C. M., Heilshorn, S. C., Blau, H. M. Bioengineering Strategies to Accelerate Stem Cell Therapeutics. Nature 2018, 557 (7705), 335–342. https://doi.org/10.1038/s41586-018-0089-z.

(6) Madl, C. M., Heilshorn, S. C. Engineering Hydrogel Microenvironments to Recapitulate the Stem Cell Niche. Annu. Rev. Biomed. Eng. 2018, 20 (1), 21–47. https://doi.org/10.1146/annurev-bioeng-062117-120954.

(7) von der Mark, K., Park, J., Bauer, S., Schmuki, P. Nanoscale Engineering of Biomimetic Surfaces: Cues from the Extracellular Matrix. Cell Tissue Res. 2010, 339 (1), 131–153. https://doi.org/10.1007/s00441-009-0896-5.

(8) Chen, L., Yan, C., Zheng, Z. Functional Polymer Surfaces for Controlling Cell Behaviors. Mater. Today 2018, 21 (1), 38–59. https://doi.org/10.1016/j.mattod.2017.07.002.

(9) Cimmino, C., Rossano, L., Netti, P. A., Ventre, M. Spatio-Temporal Control of Cell Adhesion: Toward Programmable Platforms to Manipulate Cell Functions and Fate. Front. Bioeng. Biotechnol. 2018, 6, 190. https://doi.org/10.3389/fbioe.2018.00190.

(10) Dhowre, H. S., Rajput, S., Russell, N. A., Zelzer, M. Responsive Cell–Material Interfaces. Nanomed. 2015, 10 (5), 849–871. https://doi.org/10.2217/nnm.14.222.

(11) Geiger, B., Spatz, J. P., Bershadsky, A. D. Environmental Sensing through Focal Adhesions. Nat. Rev. Mol. Cell Biol. 2009, 10 (1), 21–33. https://doi.org/10.1038/nrm2593.

(12) Guasch, J., Conings, B., Neubauer, S., Rechenmacher, F., Ende, K., Rolli, C. G., Kappel, C., Schaufler, V., Micoulet, A., Kessler, H., Boyen, H.-G., Cavalcanti-Adam, E. A., Spatz, J. P. Segregation Versus Colocalization: Orthogonally Functionalized Binary Micropatterned Substrates Regulate the Molecular Distribution in Focal Adhesions. Adv. Mater. 2015, 27 (25), 3737–3747. https://doi.org/10.1002/adma.201500900.

(13) Fumasi, F. M., Stephanopoulos, N., Holloway, J. L. Reversible Control of Biomaterial Properties for Dynamically Tuning Cell Behavior. J. Appl. Polym. Sci. 2020, 137 (25), 49058. https://doi.org/10.1002/app.49058.

(14) Yang, J., Yao, M.-H., Du, M.-S., Jin, R.-M., Zhao, D.-H., Ma, J., Ma, Z.-Y., Zhao, Y.-D., Liu, B. A Near-Infrared Light-Controlled System for Reversible Presentation of Bioactive Ligands Using Polypeptide-Engineered Functionalized Gold Nanorods. Chem. Commun. 2015, 51 (13), 2569–2572. https://doi.org/10.1039/C4CC09516B.

(15) Sur, S., Matson, J. B., Webber, M. J., Newcomb, C. J., Stupp, S. I. Photodynamic Control of Bioactivity in a Nanofiber Matrix. ACS Nano 2012, 6 (12), 10776–10785. https://doi.org/10.1021/nn304101x.

(16) Gao, T., Li, L., Wang, B., Zhi, J., Xiang, Y., Li, G. Dynamic Electrochemical Control of Cell Capture-and-Release Based on Redox-Controlled Host–Guest Interactions. Anal. Chem. 2016, 88 (20), 9996–10001. https://doi.org/10.1021/acs.analchem.6b02156.

(17) Lee, E., Luo, W., Chan, E. W. L., Yousaf, M. N. A Molecular Smart Surface for Spatio-Temporal Studies of Cell Mobility. PLOS ONE 2015, 10 (6), e0118126. https://doi.org/10.1371/journal.pone.0118126.

(18) Zhang, L., Wang, Z., Das, J., Labib, M., Ahmed, S., Sargent, E. H., Kelley, S. O. Potential-Responsive Surfaces for Manipulation of Cell Adhesion, Release, and Differentiation. Angew. Chem. Int. Ed Engl. 2019, 58 (41), 14519–14523. https://doi.org/10.1002/anie.201907817.

(19) MacCulloch, T., Buchberger, A., Stephanopoulos, N. Emerging Applications of Peptide– Oligonucleotide Conjugates: Bioactive Scaffolds, Self-Assembling Systems, and Hybrid Nanomaterials. Org. Biomol. Chem. 2019, 17 (7), 1668–1682. https://doi.org/10.1039/C8OB02436G.

(20) Freeman, R., Stephanopoulos, N., Álvarez, Z., Lewis, J. A., Sur, S., Serrano, C. M., Boekhoven, J., Lee, S. S., Stupp, S. I. Instructing Cells with Programmable Peptide DNA Hybrids. Nat. Commun. 2017, 8, 15982. https://doi.org/10.1038/ncomms15982.

(21) Zhang, D. Y., Seelig, G. Dynamic DNA Nanotechnology Using Strand-Displacement Reactions. Nat. Chem. 2011, 3, 103–113. https://doi.org/10.1038/nchem.957.

(22) Gröger, K., Gavins, G., Seitz, O. Strand Displacement in Coiled-Coil Structures: Controlled Induction and Reversal of Proximity. Angew. Chem. Int. Ed. 2017, 56 (45), 14217–14221. https://doi.org/10.1002/anie.201705339.

(23) Tripet, B., Yu, L., Bautista, D. L., Wong, W. Y., Irvin, R. T., Hodges, R. S. Engineering a de Novo-Designed Coiled-Coil Heterodimerization Domain for the Rapid Detection, Purification and Characterization of Recombinantly Expressed Peptides and Proteins. Protein Eng. Des. Sel. 1996, 9 (11), 1029–1042. https://doi.org/10.1093/protein/9.11.1029.

(24) Tashiro, K.-I., Sephel, G. C., Greatorex, D., Sasaki, M., Shirashi, N., Martin, G. R., Kleinman, H. K., Yamada, Y. The RGD Containing Site of the Mouse Laminin A Chain Is Active for Cell Attachment, Spreading, Migration and Neurite Outgrowth. J. Cell. Physiol. 1991, 146 (3), 451–459. https://doi.org/10.1002/jcp.1041460316.

(25) Kapp, T. G., Rechenmacher, F., Neubauer, S., Maltsev, O. V., Cavalcanti-Adam, E. A., Zarka, R., Reuning, U., Notni, J., Wester, H.-J., Mas-Moruno, C., Spatz, J., Geiger, B., Kessler, H. A Comprehensive Evaluation of the Activity and Selectivity Profile of Ligands for RGD-Binding Integrins. Sci. Rep. 2017, 7, 39805. https://doi.org/10.1038/srep39805.

(26) Arnold, M., Cavalcanti‐Adam, E. A., Glass, R., Blümmel, J., Eck, W., Kantlehner, M., Kessler, H., Spatz, J. P. Activation of Integrin Function by Nanopatterned Adhesive Interfaces. ChemPhysChem 2004, 5 (3), 383–388. https://doi.org/10.1002/cphc.200301014.

(27) Finke, A., Bußkamp, H., Manea, M., Marx, A. Designer Extracellular Matrix Based on DNA–Peptide Networks Generated by Polymerase Chain Reaction. Angew. Chem. Int. Ed. 2016, 55 (34), 10136–10140. https://doi.org/10.1002/anie.201604687.

(28) Zhou, D., Zhang, G., Gan, Z. C(RGDfK) Decorated Micellar Drug Delivery System for Intravesical Instilled Chemotherapy of Superficial Bladder Cancer. J. Controlled Release 2013, 169 (3), 204–210. https://doi.org/10.1016/j.jconrel.2013.01.025.

(29) Diaz, C., Neubauer, S., Rechenmacher, F., Kessler, H., Missirlis, D. Recruitment of α v β 3 Integrin to α 5 β 1 Integrin-Induced Clusters Enables Focal Adhesion Maturation and Cell Spreading. J. Cell Sci. 2020, 133 (1), jcs232702. https://doi.org/10.1242/jcs.232702.

(30) Hynes, R. O. Integrins: Bidirectional, Allosteric Signaling Machines. Cell 2002, 110 (6), 673–687. https://doi.org/10.1016/S0092-8674(02)00971-6.

(31) Lu, P., Weaver, V. M., Werb, Z. The Extracellular Matrix: A Dynamic Niche in Cancer Progression. J. Cell Biol. 2012, 196 (4), 395–406. https://doi.org/10.1083/jcb.201102147.

(32) Ellis, S. J., Tanentzapf, G. Integrin-Mediated Adhesion and Stem-Cell-Niche Interactions. Cell Tissue Res. 2010, 339 (1), 121–130. https://doi.org/10.1007/s00441-009-0828-4.

(33) Microscope Image Processing; Wu, Q., Merchant, F. A., Castleman, K. R., Eds., Elsevier/Academic Press: Amsterdam; Boston, 2008.

(34) Ronneberger, O., Fischer, P., Brox, T. U-Net: Convolutional Networks for Biomedical Image Segmentation. In Medical Image Computing and Computer-Assisted Intervention – MICCAI 2015; Navab, N., Hornegger, J., Wells, W. M., Frangi, A. F., Eds., Springer International Publishing: Cham, 2015; pp 234–241.

(35) Missirlis, D., Haraszti, T., Scheele, C. v. C., Wiegand, T., Diaz, C., Neubauer, S., Rechenmacher, F., Kessler, H., Spatz, J. P. Substrate Engagement of Integrins A5β1 and Avβ3 Is Necessary, but Not Sufficient, for High Directional Persistence in Migration on Fibronectin. Sci. Rep. 2016, 6 (1), 23258. https://doi.org/10.1038/srep23258.

